# Alanine Metabolism in *Bacillus subtilis*

**DOI:** 10.1101/562850

**Authors:** Karzan R. Sidiq, Man W. Chow, Zhao Zhao, Richard A. Daniel

## Abstract

Alanine plays a crucial role in bacterial growth and viability, the L-isomer of this amino acid is one of the building blocks for protein synthesis, and the D-isomer is incorporated into the bacterial cell wall. Despite a long history of genetic manipulation of *Bacillus subtilis* using auxotrophic markers, the genes involved in alanine metabolism have not been characterised fully. Here we identify a *B. subtilis* alanine permease, YtnA (DatA), which has a major role in the assimilation of D-alanine from the environment, and provide an explanation for the observation that growth of *B. subtilis* does not result in the significant accumulation of extracellular D-alanine. Interestingly, this transporter seems to have specificity for D-alanine but also transports L-alanine. We also show that, unlike *E. coli* where multiple enzymes have a biochemical activity that can generate alanine, in *B. subtilis* the primary synthetic enzyme for alanine is encoded by *alaT*, although a second gene, *dat*, is present that can support slow growth of an L-alanine auxotroph. However, Dat probably synthesises D-alanine and its activity is influenced by the abundance of L-alanine. Our work provide a clear insight into the complex network of alanine metabolism, and also points to the possibility that the relative abundance of D- and L-alanine might be linked with cytosolic pool of D and L-glutamate, coupling protein and cell envelope synthesis with the metabolic status of the cell.

**Author Summary:** D-alanine is a central component of the cell envelope of *Bacillus subtilis*, it has an essential role in both peptidoglycan crosslinking and is also incorporated into the teichoic acids. Growth of the bacterial cell results in the release D-alanine and as this process is on the outside of the cell it should result in the accumulation of this amino acid in the culture medium. However, this has not been see to occur if *B. subtilis* cultures. This work identifies and characterises a membrane transporter as having primary role seems to be the assimilation of D-alanine as it is released from the cell wall. The analysis shows that *in vivo* this transporter is specific for alanine, but appears to have greater affinity for the D-isomer of this amino acid. This work also clarifies the genetics of alanine biosynthesis in B. subtilis through which there are interesting difference with other bacterial species.

## Introduction

Bacterial cell growth requires the coordinated synthesis of all the metabolites necessary for the enlargement of the cell. One essential component of the bacterial cell wall that is unique to bacteria is D-alanine. This amino acid isomer is universally used by bacteria in the synthesis of the peptidoglycan (PG) as a substrate for penicillin binding protein-mediated crosslinking of the glycan strains (1, 2). In Gram-positive bacteria, it is also used to modify the teichoic acids, which are present as a second major component of the cell wall and membrane (3, 4).

Bacteria obtain D-alanine primarily by converting L-alanine by racemization. L-alanine, on the other hand, can be biosynthesised by the cell, in addition to assimilation from outside the cell and racemization of D-alanine. Alanine synthesis and metabolism is increasingly being exploited for antibacterial and industrial applications. For example, exogenous L-alanine has been shown to increase the susceptibility of some pathogens to antibiotics (5); D-alanine can inhibit germination of the spores of Bacillus and Clostridium species by acting as an competitor of L-alanine, one of the most effective germinants (6, 7); the level of D-alanine has also been shown to be able to alter the course of infection by *Bacillus anthracis* (8). In *B. subtilis* an alanine racemase gene has been utilised for plasmid maintenance (9), and further explored by Xia *et al*. (2007) to replace antibiotics resistance genes in the construction of a food-grade expression system (10).

It has long been proposed that growth of Gram-positive rod shaped bacteria results from the degradation of the outer old cell wall material (11–13) and the release of this material into the growth media. It is also known that formation of each cross-link in the peptidoglycan will result in the release of a D-alanine residue. PG peptide side chains that do not take part in cross-links are also matured soon after synthesis in B. subtilis, with both of the terminal D-alanine residues being cleaved off sequentially (14, 15). D-alanine is also released from the teichoic acids by spontaneous hydrolysis (16), consequently there should be an accumulation of free D-alanine residues in the vicinity of a growing cell unless actively re-assimilated by the bacterial cells. Pervious analyses of culture media composition after the growth of various bacterial species have revealed that many release D-amino acids on reaching stationary phase growth, presumably as secondary metabolites or signalling molecules (2). Indeed, stationary phase cultures of *S. aureus* have been shown to have significant levels of D-alanine present (1). Surprisingly, however, analysis of spent media from *B. subtilis* cultures were essentially devoid of D-alanine (1). Since both bacterial species incorporate this amino acid into the cell envelope in similar ways this suggested that *B. subtilis* has an efficient recycling system for D-alanine.

In *B. subtilis* the biosynthetic pathways for alanine are poorly characterised, and this is complicated by the partial annotation and functional characterisation of the racemase and aminotransferase genes (17). In *B. subtilis* two alanine racemases, AlrA (originally called Dal) (18) and YncD (also called Alr) (19), have been identified and it seems that YncD/Alr only functions in sporulation. *B. subtilis* also encodes two putative aminotransferases: YugH which is also called AlaT (20), and YheM (Dat) which is a homolog of the *Bacillus sphaericus* Dat protein, a D-amino acid transaminase that converts D-alanine to D-glutamate (21). Alanine degradation, on the other hand, is believed to be carried out by Ald, an alanine dehydrogenase that reversibly catalyses the reductive amination of alanine to pyruvate. Ald is required for utilisation of alanine as a sole carbon or nitrogen source (22, 23) and also for sporulation in *B. subtilis* (24). A recent characterisation of the roles of DacA/LdcB (15) in *B. subtilis* has indicated that there is a high demand for the D-isomer of alanine for cell envelope biosynthesis, as might be expected considering its presence in both peptidoglycan and teichoic acids. D-alanine can be supplied by either cytosolic isomerisation of alanine by the action of AlrA (18) or assimilation from the environment, although this isomeric form is not normally available in nature in significant concentrations. Like glutamate, alanine is abundant in the cytoplasm of growing cells, present in both the L- and D- isomeric form and there is strong evidence for the presence of mechanisms for precise control of the concentration of these amino acids and presumably their isomeric state (25). Surprisingly, in contrast to glutamate, the biosynthetic pathway for alanine is poorly defined in *B. subtilis*. Also, little is known about the uptake mechanisms for the assimilation of exogenous alanine by bacteria. Although many transporters are bioinformatically annotated as being potentially involved in alanine uptake, experimental evidence is limited and only one from *E. coli* has been implicated in D-alanine transport (26).

Here we report the identification of an alanine transporter, which we have named DatA (<u>D</u>-<u>a</u>lanine <u>t</u>ransporter <u>A</u>), that is required for efficient uptake of exogenous D-alanine. *In vivo* characterisation shows that DatA has specificity for alanine, with preference for D-alanine, and that it does not seem to function in the uptake of other amino acids. We have also clarified aspects of the synthetic pathways for both isomeric forms of alanine in *B. subtilis*. Our results provide explanations for some longstanding observations; we show that AlaT (YugH) functions as the major transaminase responsible for the biosynthesis of alanine, whereas the second enzyme Dat has a minor role in alanine biosynthesis, probably functioning in synthesis of D-alanine and its activity is inhibited by the presence of L-alanine.

## Results and Discussion

### Growth characteristics and L-alanine sensitivity of a D-alanine dependent strain of *B. subtilis*

Deletion of the alanine racemase gene, *alrA* (*dal*), has previously been shown to result in auxotrophy for D-alanine (9). We sought to use this property to determine if a specific D-alanine uptake system was present or if uptake was mediated by transporters with degenerate specificity as has been suggested for some other amino acid uptake systems (27). So we characterised the growth of a strain lacking the conditionally essential alanine racemase, *alrA* (strain RD180) in a LB growth medium. As shown by Ferrari *et al*. (9), in the absence of added D-ala the strain failed to grow in the medium, and no to poor growth was observed in the presence of less than 200 μM D-alanine. But when supplemented with D-alanine up to 0.5 mM the strain exhibited concentration dependent growth. Above 0.5 mM D-alanine the strain grew maximally irrespective of the D-alanine concentration (Fig 1A and B), showing that the strain was able to take up D-alanine efficiently enough to support growth. However, even at the highest concentration the resulting growth rate was significantly lower than that of the wild type strain (168CA) (Fig 1B), suggesting that uptake was not sufficient to supply the demands of normal growth. From these growth curves it was determined that 450 μM D-alanine was sufficient for the strain to grow optimally. We then used a fixed D-alanine concentration (450 μM) and supplemented the culture medium (LB) with different concentrations of other amino acids to determine if the presence of an excess of one particular amino acids would interfere specifically with the growth of the *alrA* null strain, and not the wild type strain 168CA. On testing all 20 amino acids it was found that only the addition of excess L-alanine resulted in any significant deviation in the growth of the *alrA* null strain but not for 168CA (Fig. 1C). None of the other amino acids tested had any significant effect on the growth of the strains and gave essentially identical growth curves to those that are shown in Fig. 1C for glycine and proline, apart from cysteine that at elevated concentrations was toxic to both strains (data not shown). This result indicated that the D-ala uptake perturbed specifically by L-alanine but no other amino acids, suggesting the existence of an alanine specific transporter and that L-alanine competitively interfered with the uptake of the D-isomer.

**Figure 1.**
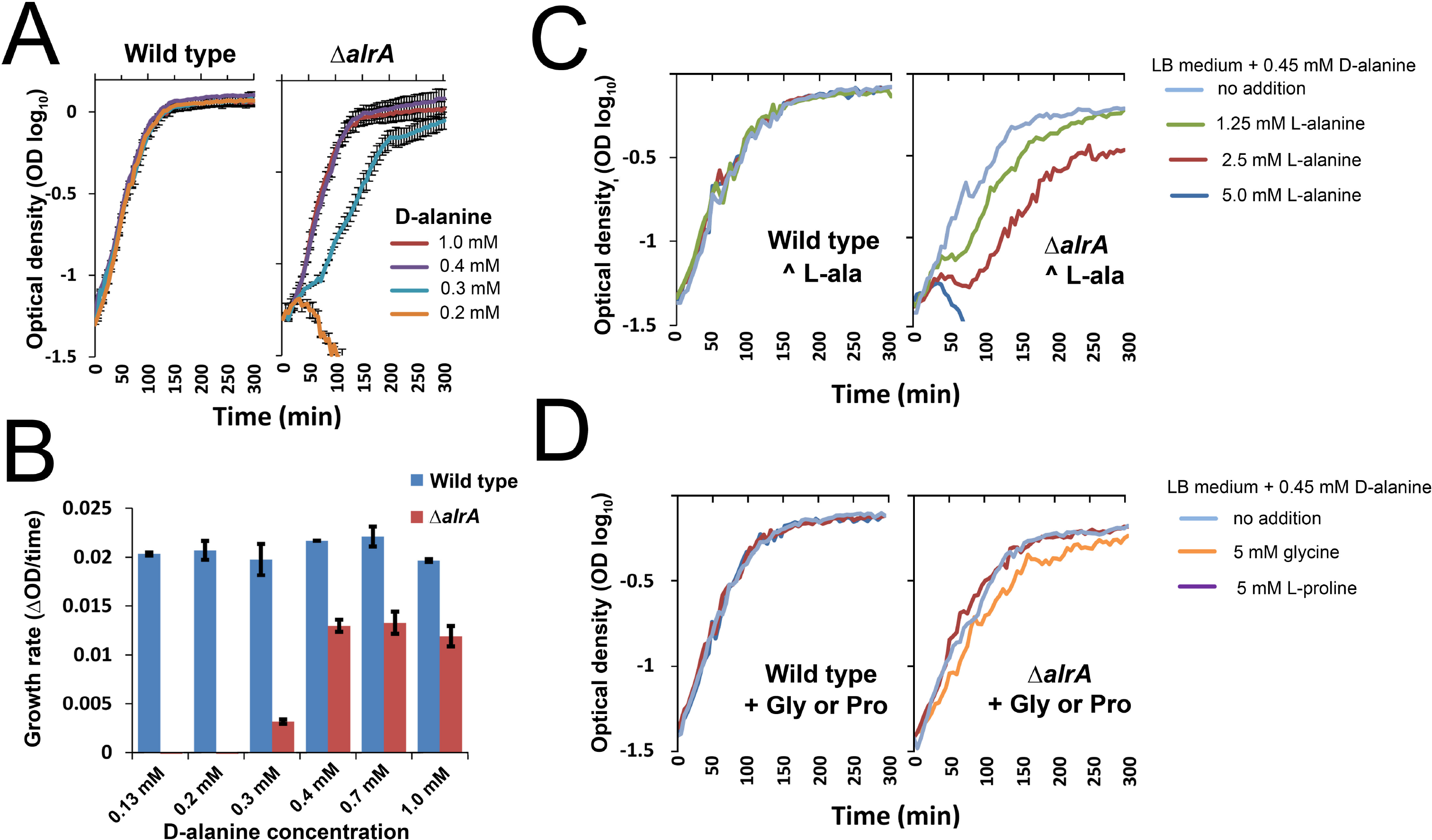
**Panel A** - The growth characteristics of D-alanine auxotroph (alrA) strain. The growth curves of wild-type (168CA) and ΔalrA::zeo (RD180) strains in LB medium, supplemented with different concentrations of D-alanine (0.2-1 mM) at 37 °C. The growth curves were monitored by plate reader and shown in Log10 of optical density (OD600). The plotted line shows the mean values for 3 biological replicates with error bars indicating the standard deviation between replicates. **Panel B** - The growth rates of the same strains measured at three time point intervals (60-80, 80-100 and 100-120 min) of panel (A). The error bars represent the standard deviation of mean of growth rate in two independent experiments. **Panels C and D** - Growth curves showing the inhibitory effect of increasing concentrations of L-alanine, glycine or L-proline on the growth of the D-alanine dependent strain (RD180) compared to the wild type strain (168CA). Both strains were grown at 37 °C in LB medium with fixed concentration of D-alanine and supplemented with different concentrations of L-alanine (C) or 5 mM Glycine or L-Proline (D) at 37 C°.

As the abundance of amino acids in LB is not well defined we used the analyses of individual media components (28) to estimate the free L-alanine concentration in LB medium to be approximately 4 mM. This was approximately 10 fold higher than the concentration of D-alanine that we used (450 μM), thus it was possible that the reduced growth rate observed of the D-alanine dependent strain (*alrA* null) was caused by the presence of excess L-alanine. To test this possibility the analysis was repeated using a defined medium (GM) where all amino acids except L-ala were present at 1 mM and D-ala was absent or present at 1 or 0.5 mM. The resulting growth of the *alrA* null strain (RD180) and the wild-type was monitored over time (Fig 2A and B). As seen previously growth of RD180, with 0.5 mM D-alanine permitting growth comparable to that seen for the wild type strain in the same medium (Fig 2A, and B). Here the slower growth of RD180 compared to the wild type was not seen, presumably because the “maximal” growth rate in this medium did not exceed the rate at which D-alanine could be taken up by the auxotroph.

**Figure 2.**
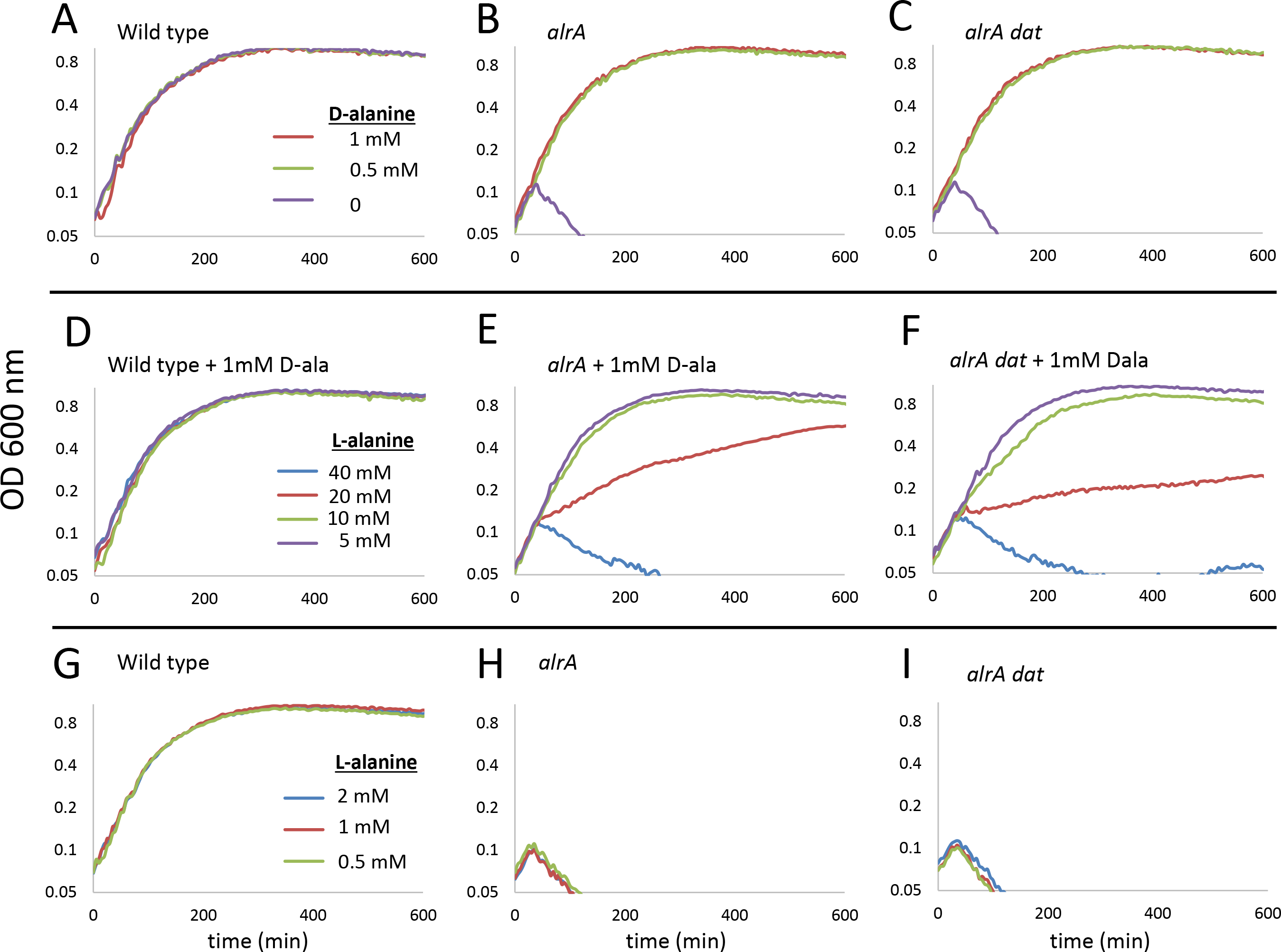
Inhibitory effect of L-alanine on the growth of a D-alanine auxotroph in a defined media. The growth of wild-type (168CA), *ΔalrA* (RD180) and *ΔalrA Δdat* (KS78) in GMaa with 1 or 0.5 mM of D-alanine or no alanine were shown in panels **A**, **B** and **C** respectively. Strains were grown in GMaa with 1 mM D-alanine supplemented with different concentrations of L-alanine (D-F), or in GMaa with L-alanine but without D-alanine (**G-I**). Growth determine in terms of optical density (600 nm) in plate reader with shaking at 37°C. Numbers in the legend represents different molarities (mM) of either D-/L-alanine (for A-C): D-alanine; for D-I: L-alanine) in the media. ‘-’ indicates no alanine was added in the media.

Using the same defined medium (GM) with D-alanine present at 1mM, the culture medium was supplemented with L-alanine at 5, 10, 20 and 40 mM and the growth of RD180 was compared with that seen for the wild type (Fig 2 D and E). Again, when the L-alanine concentration in the medium was greater than 10 times that of D-ala growth of RD180 was perturbed, and where the L-alanine was present at high concentrations (40 fold excess) the RD180 failed to grow.

It has previously been proposed that the presence of L-alanine might alter metabolism such that D-alanine is metabolised efficiently into pyruvate and so may not be available for cell wall synthesis, resulting in growth inhibition (23). However, it was equally possible that the transporter of exogenous alanine was unable to differentiate between the isomeric states of alanine and consequently, when the ratio of extracellular L- to D-alanine exceeded a certain level, D-ala uptake was insufficient to permit cell envelope synthesis (peptidoglycan and teichoic acid synthesis) due to transport competition. Strangely, however, as has been noted previously (9, 23), RD180 when inoculated on equivalent minimal solid medium was found to grow in the absence of D-alanine, albeit at a slower rate than in the presence of D-alanine (Fig 3). This indicates that under these conditions an alternate, less efficient pathway for the synthesis of D-ala was occurring. Interestingly, we found that this growth was also specifically inhibited if L-alanine is present (Fig. 3B) in the medium, suggesting that L-ala repressed the alternate D-alanine synthetic pathway.

**Figure 3.**
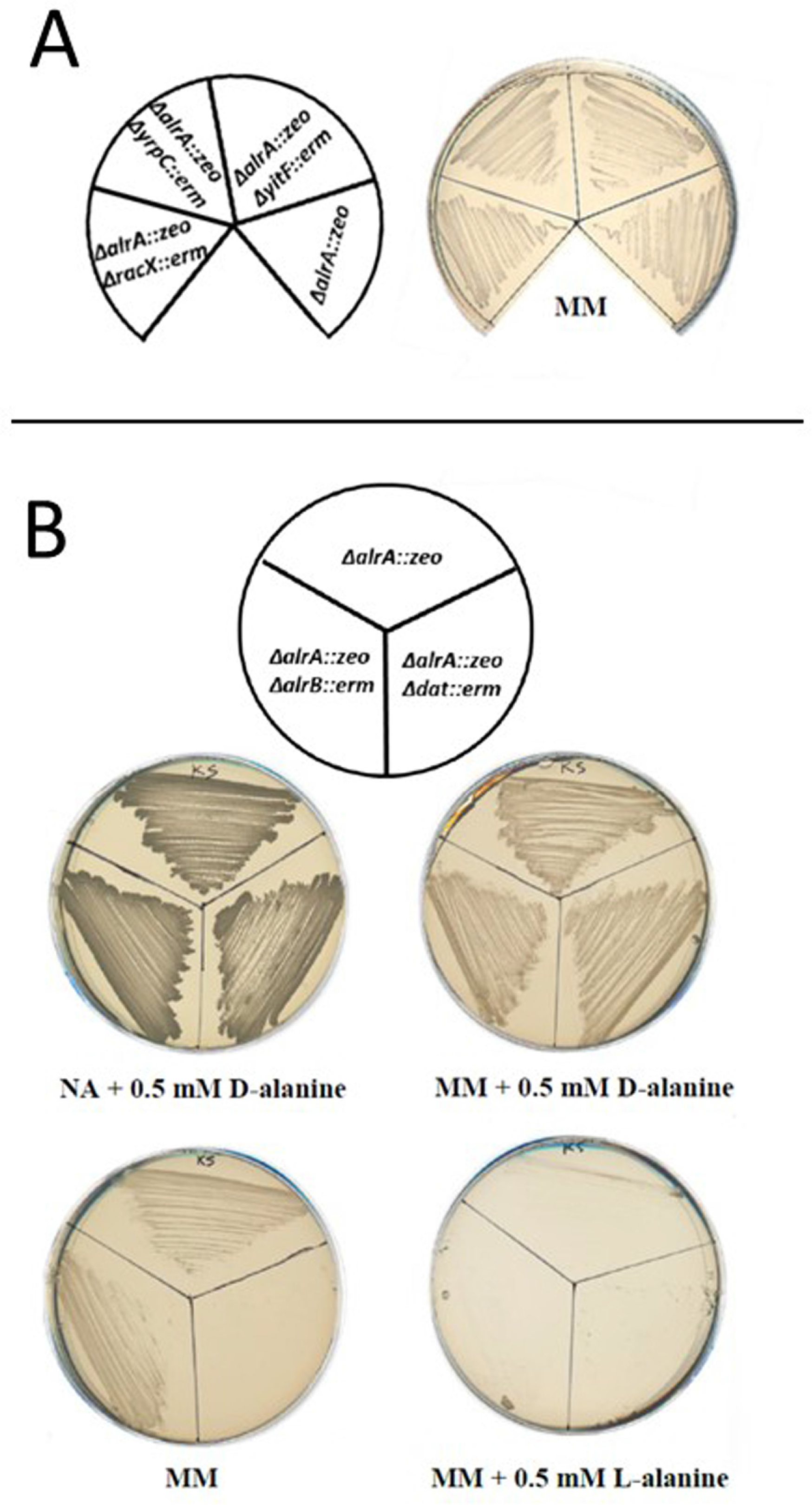
Identification of the second D-alanine synthetic enzyme. **Panel A** - The D-alanine auxotroph strains *ΔalrA* (RD180), *ΔalrA ΔyitF* (KS35), *ΔalrA ΔyrpC* (KS34), and *ΔalrA ΔracX* (KS33) were found to be able to grow well on Minimal medium (MG) in the absence of D-alanine, (plate was incubated at 37 °C for 36 hours). **Panel B** - Growth of strains *ΔalrA ΔalrB* (KS32), *ΔalrA* (RD180) and *ΔalrA Δdat* (KS78) on MM with and without supplementation of 0.5 mM D- or L-alanine, at 37 °C for 48 hours. As a positive control the strains were also inoculated on nutrient agar (NA) supplemented with D-alanine (0.5 mM)

### Identification of an alternate D-alanine synthetic pathway

To try and identify the genes involved in this second synthetic pathway for D-alanine we firstly eliminated the possibility that the sporulation racemase, *alrB* (19), might be expressed under these conditions. We introduced the *alrA* null mutation into a strain already deleted for *alrB* (Fig. 3B) and found that the *alrA alrB* double mutant was still able to grow on GM plates in the absence of D-alanine, comparable to that seen for the *alrA* single mutant (Fig. 3B). We next introduced the *alrA* null mutation into strains deleted for known non-essential racemase-like genes (*yitF* (29), yrpC (30) or *racX* (31)). Again all of these double mutants were able to grow slowly on GM plates without D-alanine (Fig. 3A). We then tested other genes known or inferred to function in alanine metabolism, including the alanine dehydrogenase *ald* (24) and the putative aminotransferase genes *yugH (alaT)* and *dat* (21), to see if they were required in a strain lacking *alrA*.

The first of these genes, *yugH*, is annotated as being similar to a PLP-dependent aspartate aminotransferase. Although it has not yet been characterised fully, it has been suggested to be involved in L-alanine biosynthesis, functioning as a transaminase and named *alaT* (A.L. Sonenshein; unpublished; (20)). Our analysis revealed that the *alaT* null mutant was auxotrophic for L-alanine and further characterisation of this gene is described later in this paper. Null mutations for either *dat* or *ald* could be easily introduced into the *alrA* mutant provided that exogenous D-alanine was present. These double mutants were then tested for the ability to grow on minimal media in the absence of D-alanine. Strain KS79, deleted for both *alrA* and *ald*, was able to grow, but a strain lacking both *alrA* and *dat* (strain KS78) was not able to propagate in the absence of D-alanine even with prolonged incubation (Fig 3B). Further analysis of the growth of this strain showed that its growth was dependent upon exogenous D-alanine under all growth conditions we have tested. Therefore, in the absence of AlrA the putative aminotransferase Dat is implicated in the synthesis of D-alanine, though at a reduced rate. Interestingly, this double mutant seemed more sensitive to the relative concentrations of D- and L-alanine in liquid culture compared to the *alrA* single mutant (Figure 2, panels C, F and I).

These results have interesting parallels with *Listeria monocytogenes* where an aminotransferase, with significant similarity to Dat of B. subtilis, has been shown to function in the synthesis of D-alanine when the supposed essential racemase was deleted (32). However, unlike *B. subtilis*, the mutant strains of *L. monocytogenes* described do not seem to exhibit any sensitivity to L-alanine. This suggested that either the expression or the biochemical activity of Dat is significantly different in this related bacterial species. This possibility is further supported by the recent characterisation of a similar enzyme in *Mycobacterium smegmatis* (33) where a Dat homologue was found to be able to compensate for null mutations is either the alanine racemase or the glutamate racemase null mutation. This property indicates that at least in *M. smegmatis* the enzymatic activity is bidirectional.

### Identification of the D-alanine uptake system

To determine if a gene encoding a D-ala specific transporter exists, a simple synthetic lethal screen was designed, using the *alrA* null mutant and systematically introducing null mutations for genes predicted to encode proteins involved in transport of solutes across the membrane. If D-alanine uptake was perturbed by the introduction of this second mutation the resulting double mutant would be inviable. In an initial screen a collection of 43 knockouts of transporter-like genes in the form of pMUTIN integrations (34) were introduced individually into the *alrA* null background, and the transformants selected for on nutrient agar with 1 mg/ml erythromycin and supplemented with 0.5 mM D-alanine. Most of those tested were found to transform into the strain efficiently; the resulting colonies had both of the expected antibiotic resistance markers and were dependent upon exogenous D-alanine. However, 3 knockout mutations, corresponding to *yfkT, ydgF and ytnA*, gave rise to very few colonies on selective plates and all of which were found to have lost the *alrA* null mutation, suggesting that the loss of either of these genes in combination with *alrA* was leathal.

To determine if these mutations were specifically perturbing the uptake of D-alanine, a second screen was developed using a strain where the *alrA* null mutation was complemented by a heterologous alanine racemase gene (*alrA* from *B. subtilis* W23 including its own promoter) cloned into the unstable plasmid, pLOSS* (35). This strain, KS21, was chromosomally deleted for *alrA* but was able to grow on normal media in the absence of exogenous D-ala, due to the racemase expressed from the *alrA*^*W23*^ gene carried by the plasmid. The plasmid (pLOSS* *alrA*^*W23*^) also constitutively expressed β-galactosidase so strains bearing the plasmid gave rise to blue colonies on media supplemented with X-gal (Fig 4A). However, pLOSS* is an unstable plasmid and can be easily lost if antibiotics selection for the plasmid is not maintained and expression of *alrA*^*W23*^ is not essential for viability. Therefore, when the media was supplemented with D-alanine the strain was able to grow with or without the plasmid, consequently white colonies would appear among blue colonies (Fig.4 B and C). But, if a mutation that interfered with D-ala uptake was present in this strain, the plasmid-borne *alrA*^*W23*^ would become essential for viability and hence only cells that carried the plasmid could grow and form blue colonies even in the presence of excess exogenous D-ala.

**Figure 4.**
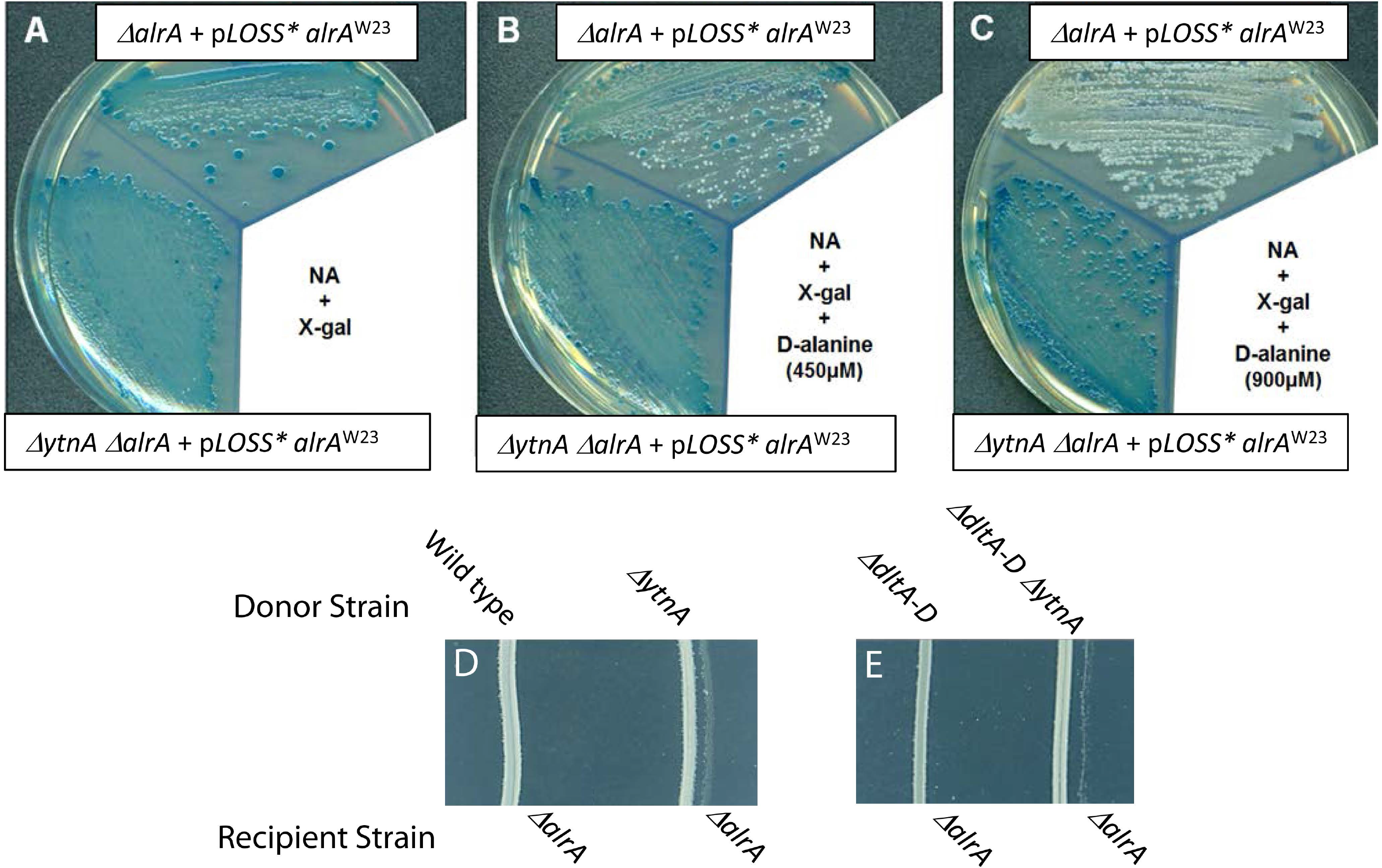
Genetic confirmation of the role of *datA* in D-alanine uptake. A D-alanine dependent strain with the sole functional copy of *alrA* encoded on a unstable plasmid was constructed, KS21 (*ΔalrA + pLOSS* Ω alrA*) and compared to a strain also lacking *datA*, KS23 (*ΔalrA ΔdatA + pLOSS* Ω alrA*) when grown on different media. **Panel A** shows the strains grown on nutrient agar (NA) plus X-gal. In contrast **panels B** and **C**show the same strains on NA + X-gal supplemented with 450 µM of D-alanine and X-gal and 900μM of D-alanine respectively. All three plates were photographed after 16 h incubation at 37 °C. The blue colonies indicate that the *pLOSS* Ω alrA* plasmid is conserved in the cells, whereas the cells produced white colonies after losing the plasmid. (D & E) In this cross-feeding assay, the 168CA, KS11 (*ΔdltA-D*), KS22 (*ΔdatA*) and KSS36 (*ΔdltA-D ΔdatA*) strains (D-alanine donors) and RD180 (*ΔalrA::zeo*) and KS12 (*ΔalrA ΔdltA-D*) strains (D-alanine recipients) were used in this assay. The D-alanine donor strains were firstly streaked on nutrient agar plates and incubated for 3 h at 37 °C. Later, the D-alanine recipient strains were streaked closely parallel to the donor strains and kept incubating for a further 23h. The total incubation time was 26 h.

The 3 mutations initially identified as potentially being synthetically lethal with *alrA* null were then introduced into the modified host strain (KS21). Transformants with both null alleles were easily obtained in all cases. However, of the 3 mutations only the deletion of *ytnA* required the presence of pLOSS\**alrA*^*W23*^ plasmid when grown in the presence of D-alanine (Fig. 4B and C). The other 2 null mutations *(yfkT* and *ydgF)* generated strains that easily lost the plasmid when D-ala was present in the culture medium, resulting in a high proportion of white colonies in the absence of spectinomycin (the resistance marker encoded on pLOSS\**alrA*^*W23*^). This indicated that these strains were able to assimilate D-ala sufficiently to allow growth without the need for the complementing copy of *alrA* on the plasmid. Thus these genetic screens identified the transporter-like protein YntA as the possible D-alanine transporter, as the loss of *yntA* resulted in a strain that was not able to exploit D-alanine in the culture medium for cell growth in the absence of the alanine racemase, AlrA.

### Cross-feeding assay

If YtnA was involved in the assimilation of exogenous D-ala, it would be expected that a strain lacking this transporter would not be able to re-assimilate D-ala released from the cell wall during growth. To test this hypothesis we employed a simple bioassay using the growth of the *alrA* null mutant as a reporter for the presence of D-alanine. Previous publications had suggested that *B. subtilis* does not release D-alanine into the culture medium during growth (1) on the basis that it was not detectable in culture supernatants. Thus the growth of a wild-type Bacillus strain was not expected to release D-ala and so would not support the growth of a D-ala auxotroph. Indeed, by streaking the *alrA* null mutant as a parallel line very close to a linear streak of the wild type strain (168CA) on a nutrient agar plate did not result in any growth of the *alrA* null mutant (Fig. 4D). In contrast, using the same inoculation technique using a *ytnA* null strain in place of 168CA resulted in clear growth of the *alrA* null strain (Fig. 4D), showing that the growth of the *ytnA* null strain resulted in the release of D-ala into the surrounding medium which could be utilised by the *alrA* null strain. In other words, D-alanine was diffusing from the *ytnA* null strain in sufficient quantity to support the growth of the D-alanine auxotroph (*alrA* null). This released D-alanine presumably originated from either the maturation of the peptidoglycan, through the action of DacA and LdcB (15), or the result of spontaneous cleavage of D-ala from the teichoic acids present in the cell envelope. Using the same cross-feeding assay, but with a strain deleted for both *ytnA* and the *dltABCD* operon (strain KS36) as the feeding strain where D-alanine is not present on the teichoic acid (3), a similar but less dramatic result was obtained (Fig. 4E), suggesting that a large proportion of released D-alanine originates from the teichoic acids under these conditions.

These results supported the idea that the putative transporter YtnA was required for D-ala uptake. We therefore rename YtnA as DatA (<u>D</u>-<u>a</u>lanine <u>t</u>ransporter <u>A</u>). The above results also show that the most likely sources of exogenous D-alanine in vegetative growth are from the teichoic acids and the maturation of the cell wall. It also demonstrated that the wild-type strain efficiently assimilated the released D-alanine, presumably for reuse in future cell wall synthesis.

### Detection of alanine in liquid cultures

Since the D-alanine released by the *datA* null strain when grown on agar plates was sufficient to support the growth of the D-alanine auxotroph, it was expected that in liquid growth the released D-ala would be detectable at the chemical level. For this parallel cultures of 168CA and KS22 (*datA* null) were grown in LB medium and the culture supernatants were derivatised with Marfeys reagent prior to HPLC analysis (36) to permit separation and quantitation of optical isomers of alanine. Using a dilution series of D-ala in sterile LB medium as standards it was possible to identify the peak corresponding to D-ala on the HPLC trace (Fig S1) and generate a standard curve for the quantitative determination of D-ala in the sample. Using this standard curve we could show that D-ala was present in the culture medium of *B. subtilis* during exponential growth (Fig S2A and B). As expected, for the wild type strain the concentration was low, and reduced to undetectable levels at the end of exponential growth. In contrast, the *datA* null strain exhibited elevated levels of D-ala throughout exponential growth (Fig.S2B). A similar result was obtained for culture supernatants of the *dltABCD* null strains, in which D-alanine was not present on the teichoic acids (Fig. S2C), although the amount of released D-alanine was less, representing only that derived from the peptidoglycan. These results were consistent with the cross feeding results on plates.

To further clarify the role of DatA in alanine uptake/release the wild-type and the *datA* null strain were grown in a defined minimal media supplemented with either 0.5 mM L-alanine, D-alanine or in the absence of any alanine. The growth of the culture was then monitored and samples were taken at regular intervals to allow the concentration of free alanine in the culture medium to be determined using derivatisation with OPA and HPLC analysis (see methods). The results obtained were then plotted against time and are shown Fig 5 (which shows a characteristic experiment). When the strains were grown in the absence of alanine the HPLC analysis clearly showed a peak corresponding to alanine appearing after about 90 min of growth. However, using OPA labelling technique the isomeric form of alanine cannot be determined. As with the LB medium (Fig S2A and B), for the wild-type strain this peak was relatively small, corresponding to about 5-7 μM and was seen to disappear toward the end of exponential growth (Fig. 5A). In dramatic contrast a significantly stronger alanine peak was obtained for the *datA* null strain starting at about the same time, but rapidly increasing to over 100 mM and remaining at this level. Where alanine was present in the culture medium it was clear that the wild-type strain efficiently assimilated this amino acid in both isomeric forms over the duration of the experiment (Fig 5B), reducing its concentration to undetectable levels between 120 and 150 min of incubation, whereas the *datA* null strain was unable to efficiently utilise either isomeric form of alanine.

**Figure 5.**
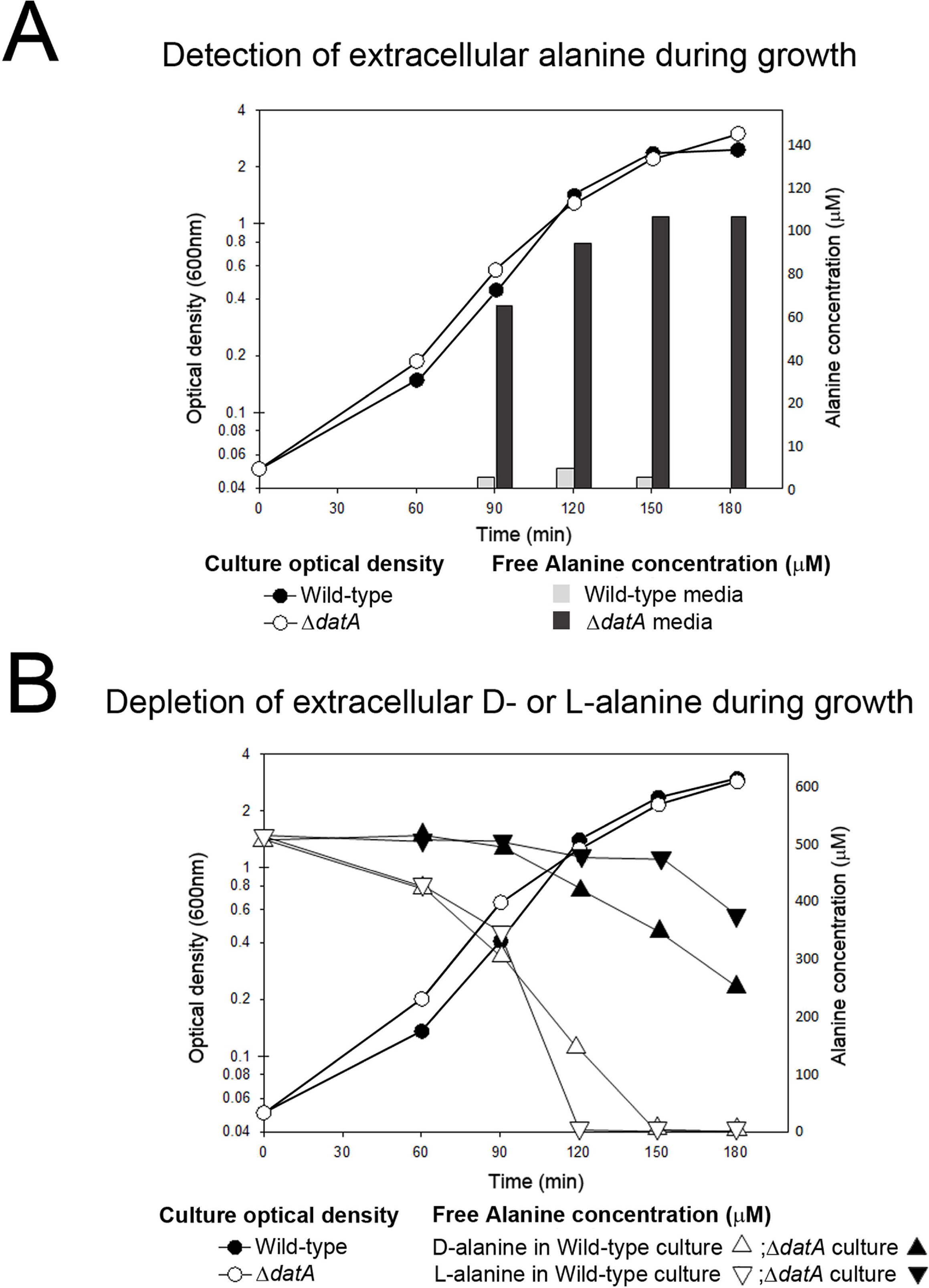
Determination the concentration of alanine present in the culture medium during growth of wild-type *B. subtilis* and a *datA* null strain by OPA derivatisation and HPLC. The line graphs show growth curves expressed as the optical density of the cultures at 600 nm (○ symbols for the wild type and • for the *datA* null strain and the column plots in panel **A** indicate the concentration of free alanine (μM). Panel B presents the growth curves for the wild-type strain and the *datA* null strain when grown the presence of exogenous alanine (0.5 mM), for simplicity the growth curves shown are for the cultures supplemented with D-alanine as those supplemented with L-alanine were essentially identical. On the same plot the alanine concentration in the respective cultures was determined over time with triangles for L-alanine (Δ for the wild-type culture and ▴for the *datA* mutant) and inverted triangles for D-alanine (▽ for the wild-type and ▾ *datA* mutant).

Interestingly, we reproducibly observed that the wild-type strain more efficiently utilised the D-isomeric form of alanine. In contrast, the poor uptake of alanine exhibited by the *datA* mutant was more evident for L-alanine than for the D-isomer. We also found that, of the other amino acids tested that exhibited depletion over time in the culture medium of the wild-type strain, only alanine was reproducibly and significantly changed by the deletion of *datA* (data not shown).

From the chemical analysis and the cross feeding experiments, it is clear that alanine is released as *B. subtilis* grows, and strains lacking DatA do not seem as efficient as the wild type strain in the re-assimilation of this amino acid. It was however evident that the concentration of alanine reduced at the end of exponential growth (Fig. 5B), suggesting that there may be other uptake systems acting to assimilate alanine in later phases of growth. However, it would seem that these are not sufficient to support vegetative growth as indicated by the synthetic lethality of *datA* and *alrA* mutations.

### Genetic context and expression of *datA*

The published transcriptional data (37) suggested that *datA* was the last gene in an operon with transcription starting upstream of *metK* and passing through *asnB* before reaching *datA*. To confirm this we designed a large set of oligonucleotides that would permit the amplification of various regions surrounding *datA* using RtPCR (Fig S3B). The functionality of these primer pairs were first confirmed using genomic DNA extracted from 168CA as a template (Fig S3C). These oligonucleotide combinations were then used to generate cDNA from total RNA extracted from exponentially growing 168CA (Fig S3D). The resulting DNA fragments were then resolved on an agarose gel to provide an indication of the size of the possible RNA transcripts in this region of the genome. This analysis supports the previous microarray data (37) suggesting that *datA* is co-transcribed with *asnB* and *metK*, both of which have been shown to be required for growth ((38) and (39) respectively).

Although *datA* is the last gene in the operon, we thought it necessary to exclude the possibility that the phenotypic effects we observed for the insertional deletion of *datA* could be due to some unforeseen effects on the expression of *asnB* and/or *metK*. For this we tested whether the phenotype could be fully complemented by ectopically expressed *datA*. Initially this was tested by introducing into the *alrA* null mutant (KS27) a copy of *datA*, using its own ribosome binding site (RBS) under the control of the IPTG-inducible P_*spac*_ promoter into the *amyE* locus. The resulting strain then transformed with the *datA* null mutation using chromosomal DNA from strain BKE30530 (*ΔdatA::erm*). However, none of the resulting colonies carried the correct markers, suggesting that the construct at *amyE* was not functioning. Examining the putative RBS of *datA* raised the possibility that translation of the ectopic copy of *datA* was poor. Thus, the inducible construct was reconstructed, this time changing the RBS to one closer to the consensus. A strain carrying this new construction as well as the *datA* deletion (KS41) was found to be easily transformed with the *alrA* null mutation in the presence of the inducer (0.1 mM IPTG; Fig S3A). Thus, enhancing the translation of the *datA* transcript was sufficient to complement the chromosomal null mutation, showing that the observed phenotype was caused solely by the deletion of *datA* and not due to negative effects on the genes upstream, and that efficient expression of native *datA* seems to be coupled with the translation of the upstream genes.

### Characterisation of an L-alanine auxotroph

As mentioned above, we found that deletion of the putative alanine dehydrogenase gene, *alaT*, led to a severe growth defect on glucose minimal media without alanine (Fig. 6A). The defect was corrected by the addition of either D- or L-alanine in the growth medium (Fig. 6A and B), suggesting that AlaT played a major role in the biosynthesis of alanine. When incubated for an extended time this null mutant was able to generate visible biomass in the absence of alanine (Fig. 6A; top right quadrant), suggesting the existence of another, inefficient biosynthetic pathway. This result is consistent with an unpublished observation by Sonenshine and Belitsky (20), that deletion of *alaT* caused severe, but not complete alanine auxotrophy. It was possible that some other transaminases were able to catalyse the necessary conversion of precursors into alanine, as has been demonstrated for *E. coli* (40). However, as the growth of the strain was very slow in the absence of alanine (Fig. 6A), we thought it also possible that Dat, the putative aminotransferase that we identified as the “secondary” D-ala synthetic pathway, has generated limited amount of D-ala which was then converted to L-ala by the racemase, to permit limited growth (Fig. 6A). To test this we constructed a strain deleted for both *alaT* and *dat* (strain MC4). Indeed, MC4 was unable to grow on glucose minimal plates without alanine. These results show that AlaT is the main enzyme for L-alanine synthesis while *dat* has a minor role. Interestingly, although expression of *dat* has been shown to be upregulated in response to nitrogen starvation (41), we found no evidence of Dat being involved in the utilisation of D-or L-alanine as a carbon or ammonium source. Instead, this role was specifically fulfilled by the alanine dehydrogenase encoded by *ald* (Fig. S4) as had previously been suggested (42).

**Figure 6.**
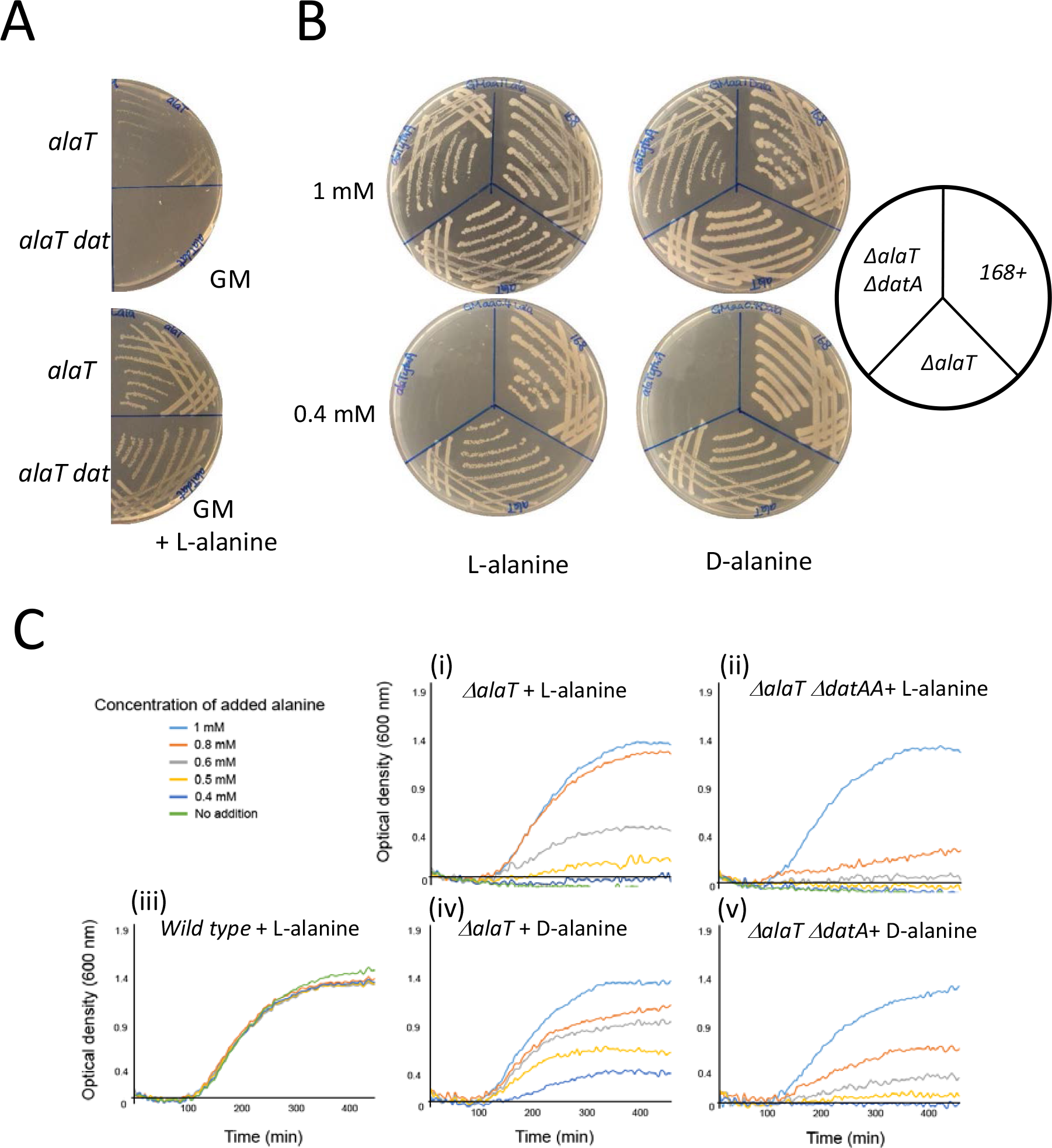
DatA transports both isoforms of alanine. **Panel A** shows how the growth of Δ*alaT* (MC2) and Δ*alaT* Δ*dat* (MC4) on GM plates is altered by the presence or absence of 1mM L-alanine. The plates were incubated at 37°C for 48 hours. **Panel B** Shows how the growth of wild-type (168+), Δ*alaT* (MC2) and Δ*alaT* Δ*datA* (MC8) on GMaa plates with either D-/L-alanine at higher (1 mM) and lower (0.4 mM) concentrations. The plates were incubated at 37°C for 24 hours. **Panel C** shows typical growth curves obtained for the wild-type strain (168), the *alaT* null strain (MC2) and the double mutant *alaT datA* (MC4, identical results were obtained for strain MC8) grown at 37 °C in a plate reader in the liquid GMaa supplemented with a range of concentrations of D-/L-alanine from 0.4 mM to 1 mM (C).

### DatA can transport both D and L-alanine

Our phenotypic characterisation of the *datA* null mutant clearly identified DatA as the transporter of D-ala. However, in the initial analysis of the *alrA* null strain we observed that excess L-ala in the culture medium interfered with the strain’s ability to grow. An *alrA dat* double mutant that absolutely depended upon exogenous D-alanine under all growth conditions tested exhibited even higher sensitivity to the level of L-alanine. This suggested that L-alanine competed with D-alanine for the transporter, raising the possibility that DatA could also transport L-alanine and was not specific for the D-isomer. This possibility was further supported by the fact that both D- and L-alanine assimilation was perturbed in the *datA* null strain (Fig. 5B). But as the strains used in these analyses were still capable of alanine biosynthesis, alanine uptake was not necessary for growth, and so might not represent the uptake capability of an alanine auxotroph. With AlaT being identified as the major biosynthetic enzyme in the alanine pathway, it was possible to determine if DatA was also required for L-alanine uptake. We constructed a strain deleted for both *alaT* and *datA*, and then tried to propagate the strain in/on GM media supplemented with L-alanine. Initial results suggested that DatA was not required for L-alanine uptake as the double mutant appeared to grow normally on minimal media supplemented with 1 mM L-alanine (Fig.6B, upper plates). However, on plates where L-alanine was provided at a lower concentration (0.4 mM) the double mutant failed to show any growth (Fig 6B, lower plates). To confirm these observations in a more quantitative manner we analysed the growth of the strain in liquid culture using a range of alanine concentrations (Fig 6C). From the growth curves generated it was clear that normal growth of an alanine auxotroph required the presence of at least 0.8 mM alanine in the culture medium (Fig. 6C (i)), whereas the *alaT datA* double mutant required at least 1 mM to exhibit the same growth profile (Fig. 6C (ii)). Similar results were also obtained when D-alanine was used to feed the strain (Fig. 6 (iv) and (v)). Surprisingly, D-ala at concentrations over 1 mM were able to support the growth of an *alaT* alanine auxotroph even when *datA* was deleted, suggesting that both D and L-alanine were able to be taken up by some other transport systems, but only when present in high concentrations.

## Summary

D-alanine is an essential component of all bacterial cell walls but it is unclear what happens to old wall as the cell grows. For Gram-negative bacteria, it is logical that the degraded material is contained within the space between the outer and inner membranes, and is available for recycling. But for Gram-positive bacteria it has been assumed that the old wall is lost to the extracellular environment. This work indicates that for D-alanine there is an efficient system for recycling of this amino acid in *B. subtilis* and that this system is probably conserved in other Bacillus spp. Our work also suggests a link between extracellular cell wall maturation and intracellular metabolism in a way that maximises the use of metabolites that would otherwise be lost to the environment, comparable to the recovery of cell wall sugars (43, 44). The results also provide an insight into the complexity of cellular metabolism, in terms of how biosynthetic pathways may be interlinked in such a way as to compensate for deficiencies in specific precursors. This analysis initially focused specifically on the uptake of a specific, non-canonical amino acid, D-alanine and exploited the predicted synthetic lethality expected for a D-alanine auxotroph that lost the ability to assimilate exogenous D-alanine. We identified a potential permease gene, *ytnA*, as a transporter gene that is necessary for efficient D-alanine uptake. Consequently, we have changed the name of *ytnA* to **<u>D</u>**-**<u>a</u>**lanine **<u>t</u>**ransporter **A**, *datA*, to reflect its apparent function in the recycle of the D-alanine that is released during the normal processes of cell wall synthesis and maturation. This role seems consistent with growth in a nutrient limited environment. Genes with significant similarity to *datA* can be identified in most *Bacilli* (not shown), although none have been characterised. Interestingly, *S. aureus* is known to release D-alanine at the end of exponential growth, at least in laboratory culture (1), suggesting that *S. aureus* does not possess this recycling function. However, genes encoding homologues of DatA can be tentatively identified by similarity, but since the similarity is low (below 48 % identity), they may have other biological roles.

Characterisation of the phenotype of the D-alanine auxotroph shows that the uptake of exogenous D-alanine is not sufficient to support high growth rates seen for a wild-type strain even in the presence of excess D-alanine (Fig. 1), indicating that the uptake is the rate limiting step. Unfortunately attempts to biochemically characterise DatA were not successful as we were unable to purify the protein. Thus we employed phenotypic and genetic methods to try and determine the specificity of the transporter. To make this possible we defined the late biosynthetic steps of L-alanine synthesis in *B. subtilis* and found that, in contrast to *E. coli* where a functionally redundant set of 3 transaminases (AvtA, YfbQ and SerC; (40)) participate in alanine synthesis, in *B. subtilis* alanine synthesis seems to be primarily the role of the transaminase, AlaT. Surprisingly, in our analysis of the D- and L- alanine dependent strains (*alrA* and *alaT* mutants), a second synthetic pathway for alanine was identified involving the D-alanine transaminase, Dat, though this only supported limited growth in both auxotrophs. The availability of an alanine auxotroph then allowed us to show that DatA is required for efficient uptake of exogenous L-alanine only when the amino acid is present at concentrations below 1 mM. This indicates that L-alanine uptake can occur through other non-specific transporters, but these have low affinity and so do not function well when the substrate is present a low concentrations. However, since this is an *in vivo* assay other factors may also be influencing uptake, such a growth of the strain that we cannot take into account.

In the light of these results we were prompted to re-evaluate alanine metabolism in *B. subtilis*. Previous analysis of other enzymes acting in alanine metabolism is rather limited, with two enzymes being identified in the literature from various Bacillus species. The first being an alanine dehydrogenase (Ald (42)) that had previous been identified as functioning in the use of alanine as a carbon source in sporulation; the second a D-alanine transaminase (Dat (45)) proposed to function in the metabolic conversion of D-alanine into D-glutamate (46), perhaps in stationary phase growth or during sporulation. However, transcriptional micro-array data and proteomics for both of these genes seems to suggest that these enzymes are maintained at a relatively constant abundance in cells (41), inferring more of a central role in cellular metabolism. Our analysis confirms that Ald is required for alanine to be used as a carbon source (Fig. S4), with the suggestion that this enzyme acts on only the L-isomer and Dat is implicated in the synthesis of both D- and L-alanine, either directly or in combination with the racemase (AlrA). Interestingly, the activity of Dat is inefficient in *B. subtilis* and is unable to fully correct the auxotrophy caused by the loss of AlaT (the main L-alanine synthase) or AlrA (the alanine racemase for D-alanine production). It was also evident that exogenous L-alanine inhibits growth of a strain lacking the alanine racemase, suggesting that the activity of Dat is modulated in some way by the availability of the L-isomer of alanine. These results together indicate that alanine biosynthesis could be integrated with the L-/D- glutamate biosynthetic pathway via Dat in such a way as to permit the generation of D-alanine, which in turn can be converted into the L-isomeric form for protein. This integration of pathways seems to be supported by the characterisation of other bacterial species, where it has been shown that Listeria possess an equivalent Dat enzyme which seems to be fully capable of supplying the cells D-alanine in the absence of a functional alanine racemase (32), though the mutant strains do not exhibit L-alanine sensitivity. This idea is further supported by the characterisation of an enzyme comparable to Dat in *Mycobacterium smegmatis* showing that it can supply the cell with D-glutamate in the absence of the cognate racemase, MurI, as well as provide D-alanine in a strain deleted for the alanine racemase (33). Thus, it would seem this set of enzymes are well conserved in a diverse range of bacteria, but most obviously in Firmicutes and close relatives. However, their activity must be modulated in different ways, presumably reflecting different metabolic priorities. This then raises the question of how the activity of these enzymes are modulated in the cytosol to ensure that the necessary precursors are available and that the metabolic pathways do not undergo futile cycles of interconversion. Our work is now focused on the subcellular localisation of these enzymes in combination with determining biochemical activity and substrate preference.

### General Material and methods

*Escherichia coli DH5α* was used for plasmids amplification. *Bacillus subtilis* strains were mutant derivatives of *168CA* and their relevant characteristics are shown in Table 1. All the chemicals and reagents used in this work were obtained from Sigma except where stated otherwise.

**Table 1.** Strains collection.

| *B. subtilis* strains | Genotype | Construction or Source or Reference |
| --- | --- | --- |
| 168 CA | *trpC2 Bacillus subtilis* | Laboratory strains collection |
| 168 + | Prototrophic derivative of 168 CA | R. Daniel, unpublished |
| W23 | *Bacillus subtilis* W23 | Laboratory strain collection |
| RD180 | *trpC2 ΔalrA::zeo* | R. Daniel, unpublished |
| BKE30530 | *trpC2 ΔdatA::erm* | BGSC (51) |
| BKE 17640 | *trpC2 ΔalrB::erm* | “ |
| BKE 34430 | *trpC2 ΔracX::erm* | “ |
| BKE09670 | *trpC2 Δdat::erm* | “ |
| BKE31400 | *trpC2 ΔalaT::erm* | “ |
| BKE31930 | *trpC2 Δald::erm* | “ |
| BKE 26810 | *trpC2 ΔyrpC::erm* | “ |
| BKE10970 | *trpC2 ΔyitF::erm* | “ |
| KS11 | *trpC2 ΔdltABCD::cat* | deletion of *dltABCD* genes |
| KS21 | *trpC2 ΔalrA::zeo pLOSS\*alrA* | *pLOSS\*alrA* transformed into RD180 |
| KS22 | *trpC2 ΔdatA::erm* | 168CA transformed with gDNA of BKE30530 |
| KS23 | *trpC2 ΔdatA::erm ΔalrA::zeo pLOSS\*alrA* | KS21 transformed with gDNA of BKE30530 |
| KS26 | *trpC2 ΔdatA::erm ΔamyE (cat P<sub>spac</sub> datA)* | pKS1 transformed into KS22 |
| KS27 | *trpC2 ΔalrA::zeo ΔamyE (cat P<sub>spac</sub> datA)* | pKS1 transformed into RD180 |
| KS32 | *trpC2 ΔalrA::zeo ΔalrB::erm* | RD180 transformed with gDNA of BKE17640 |
| KS33 | *trpC2 ΔalrA::zeo ΔracX::erm* | RD180 transformed with gDNA of BKE34430 |
| KS34 | *trpC2 ΔalrA::zeo ΔyrpC::erm* | RD180 transformed with gDNA of BKE 26810 |
| KS35 | *trpC2 ΔalrA::zeo ΔyitF::erm* | RD180 transformed with gDNA of BKE10970 |
| KS36 | *trpC2 ΔdatA::erm ΔdltA-D::cat* | KS22 transformed with gDNA of KS11 |
| KS41 | *trpC2 ΔdatA::erm ΔamyE(cat P<sub>spac</sub> ftsL(RBS)- datA )* | pKS6 transformed into KS22 |
| KS42 | *trpC2 ΔdatA::erm ΔamyE (cat P<sub>spac</sub> ftsL(RBS)- datA) ΔalrA::zeo* | KS41 transformed with gDNA of RD180 |
| KS78 | *trpC2 ΔalrA::zeo Δdat::erm* | RD180 transformed with gDNA of BKE09670 |
| KS79 | *trpC2 ΔalrA::zeo Δald::erm* | RD180 transformed with gDNA of BKE31390 |
| MC1 | *ΔalaT::erm* | gDNA of BKE31400 transformed into 168+. |
| MC2 | *ΔalaT* | Erythromycin marker was removed using pDR244. |
| MC3 | *Δdat::erm* | gDNA of BKE09670 transformed into 168+. |
| MC4 | *ΔalaT Δdat::erm* | gDNA of BKE 09670 transformed into MC2. |
| MC5 | *Δald::erm* | gDNA of BKE 09670 transformed into 168+. |
| MC6 | *ΔalaT Δald::erm* | gDNA of BKE 09670 transformed into MC2 |
| MC7 | *ΔdatA::erm* | gDNA of BKE 09670 transformed into 168+. |
| MC8 | *ΔalaT ΔdatA::erm* | gDNA of BKE 09670 transformed into MC2. |
| ***E. coli* strains** | **Genotype/Relevant features** | **Construction or Source or Reference** |
| DH5α | *fhuA2 Δ(argF-lacZ)U169 phoA glnV44 Φ80 Δ(lacZ)M15 gyrA96 recA1 relA1 endA1 thi-1 hsdR17* | New England BioLabs® |

Plasmid DNA was routinely extracted using Plasmid MiniPrep Kit (Qiagen), whilst the extraction of *B. subtilis* chromosomal DNA was either done using DNA wizard kit (Promega) for sequencing and PCR, or as a simple DNA extract (47) to permit strain construction. The transformation of *B. Subtilis* strains was carried out according to the method developed by Anagnostopoulos and Spizizen (48), modified by Young and Spizizen (49) and *E. coli* transformation was done as described by Hanahan,1985 (50). Selection for antibiotic resistance markers was done using the following concentrations: chloramphenicol (5 μg/ml), erythromycin (1 μg/ml), spectinomycin and ampicillin (100 μg/ml) in Nutrient Agar (Oxoid). Where required 5-bromo-4-chloro-3-indolyl-β-D-galactopyranoside (X-gal) was added to a final concentration of 0.002% to provide a visual indication of β-galactosidase activity, and where required IPTG was added to a final concentration of 1 mM.

### Bacterial strains and culture manipulation

The bacterial strains were routinely grown in LB (Lysogeny broth) and on nutrient agar with added supplements when required (0.5 mM D-alanine; 1 mM IPTG). Where bacterial strains were grown in a defined media based on Spizizen minimal medium (SMM, Anagnostopoulos and Spizizen (48), Young and Spizizen (49); 0.2 % w/v ammonium sulphate, 1.4 % dipotassium phosphate, 0.6 % potassium dihydrogen phosphate, 0.1 % sodium citrate dehydrate, 0.02 % magnesium sulphate, to which was added 5 mM magnesium sulphate heptahydrate, 0.1 mM calcium chloride dehydrate, 65 μM manganese (II) sulphate tetrahydrate) this provided a basic salts medium to which supplements were added to permit growth (where necessary glucose was added to a final concentration of 10g/l and each amino acids were present at a final concentration of 1.25 mM each except where specifically stated). However where amino acid utilisation as a nitrogen source was investigated the SMM was modified by omitting the ammonium sulphate. To simplify the description of the media we have adopted M to denote minimal with any omissions in brackets followed by the additions, G for glucose, A for amino acids or the three letter code for individual amino acids, again indicating any omitted components in brackets. Hence, MGA indicates the medium with salts, glucose and all amino acids added, while M-NH3 G D-Ala indicates that the minimal medium has the salts, but lacks the ammonium sulphate, Glucose and D-alanine.

For growth curve assays in LB, the bacterial strains of interest were grown in LB or defined media (supplemented with ~450 μM of D-alanine where necessary) at 30°C overnight. The overnight cultures were diluted 1:30 with pre-warmed medium and grown for about 2 hours at 37°C to obtain exponentially growing cultures. The optical density of the culture were determined (OD_600nm_) and measured volumes of the cultures were centrifuged for 3 minutes in a benchtop centrifuge (RCF 9,509). The cell pellets were quickly suspended in pre-warmed culture medium and dispensed into pre-warmed flasks or the wells of microplates, which were already loaded with pre-warmed medium such that the final OD_600_ in each flask was ~ 0.1 in each flask and ~ 0.05 in each well of microplate. The cultures were then incubated with shaking and the optical density (OD_600nm_) measured at regular intervals using a spectrophotometer or automatically by the plate reader (BMG Fluorostar). Where the experiment required the medium composition to be changed cultures were grown overnight in a complete medium and back diluted as described above into the same medium and incubated for 2 h. The optical density at 600 nm (OD_600_) was the determine and measured volumes of the cultures were centrifuged at 13,300 rpm for 3 minutes in a benchtop centrifuge (Heraeus Pico17, 45° angle rotor, 24 places, RCF 17000 × g). The pellets were suspended in appropriate volumes of the new medium and distributed into culture vessels, either Flasks or sterile microplates that were pre-loaded with appropriate volume of media with relevant supplements such that the correct final optical density was obtained. The growth of the strains was then monitored as described above.

### Determination of growth on solid media

For growth analysis using solid media, agar plates (1.5 % w/v) were prepared using SMM salts to which was added 5 mM magnesium sulphate, 0.1 mM calcium chloride dehydrate and 65 μM manganese (II) sulphate. Variations of this base medium were made by the addition of different supplements: in general 10 g/l glucose was added as a carbon source (Glucose minimal; GM), plates supplemented with 98 μM L-tryptophan (GMT) or 1 mM L-alanine; GMTala and GMala plates were supplemented with L-tryptophan and 1 mM D-alanine or alanine alone, GMaa plates contained 1.25 mM of the 19 standard amino acids omitting alanine. For plates testing for utilisation of alanine as sole carbon source, glucose was removed from SMM and replaced with 25 mM L- or D-alanine (Mala). For those testing for utilisation of alanine as a sole nitrogen source, ammonium sulphate were removed from SMM salt mix and the medium supplemented with 10 g/l glucose and 25 mM alanine (GM-NH_3_ ala plates). Strains were streaked on prepared agar plates and incubated at 37°C for 24 hours (GMaa plates) or 48 hours (GMT, GM, GMala, Mala and GM-NH_3_ala plates) and growth determined by visual inspection was recorded by photography.

### Bacterial strain construction

Mutants with single gene deletion were obtained from the Bacillus Genetic Stock Centre (BGSC) where single gene deletions had been generated by insertional deletion using a erythromycin resistance cassette (51). Chromosomal DNA from these strains at low DNA concentrations were used to transform the wild-type strains 168CA or 168+ to ensure isogenic strains were characterised. Where necessary the temperature-sensitive vector pDR244 (51) with a functional *cre* recombinase gene was used to loop out the erythromycin marker between the lox sites, leaving the resultant strain a marker-less deletion mutant. Repetition of this method was used to generate double or triple mutants where required. The transformed strains were checked that they had the appropriate antibiotic resistance, and the gene knockouts were confirmed by PCR. The list of constructed strains of *B. subtilis* is shown in Table 1.

To identify potential D-alanine synthesis pathways, searches of the bioinformatics databases (*Subti*Wiki, SubCyc and Metacyc) were used to identify any genes that have either putative or been functionally characterised to have roles in D-amino acid metabolism. Strains deleted for the genes identified were obtained from BGSC and are listed in Table S4.

The deletion of the *dlt* operon was generated by ligation of 3 DNA fragments generated by PCR. Firstly, oligonucleotides oKS01 and oKS02 were used to amplify a 2.0 kb DNA fragment, complementary to the DNA sequence upstream of the *dltA* gene. Then oKS03 and oKS04 were used to generate a 2.0 kb DNA fragment at the downstream of the *dltD* gene. Both of these DNA fragments were then digested with *Bg/II*, and after purification, appropriate quantities of the digested DNA fragments were ligated with a chloramphenicol resistant cassette (cat) obtained from plasmid pSG1 as a *BamH*I fragment. The ligation product was used directly in the transformation of *B. subtilis* 168CA strain. The correct knockout strain (Δ*dltABCD∷cat*; KS11) was isolated as a chloramphenicol resistant transformant and confirmed by PCR.

### Amino acid competition assay

The wild-type strain (168CA) and the D-alanine dependent strain RD 180 (*ΔalrA::zeo*) were grown in LB medium, supplemented with 450 μM of D-alanine to which other individual amino acids (Melford Bioscience) at concentrations between 0.0098 mM – 5.0 mM were added. The growth of the strains at 37 °C was then monitored over time using microplate reader (BMG, Fluorostar). The readings obtained were then used to plot growth curves for the culture to determine if the addition of a specific amino acid had any specific significant impact on D-alanine dependent strain (RD180) and not on the wild-type strain (168CA).

### Synthetic lethal screen

To determine if a specific transporter was required for the uptake of D-alanine by the D-alanine auxotroph, (strain RD180) the auxotroph was systematically transformed with insertional mutations in genes potentially encoding amino acid transporter proteins. Using the premise that the loss of this transporter in the *alrA* null background would be lethal and very few colonies bearing the incoming resistance marker would be obtained compared to other null mutations. Our initial screen utilised the pMUTIN knockout collection (obtained from the NBRP strain collection, Japan (52)) as a source of null mutations and a list of the mutations tested is shown in Table S4. Where a significant reduction in transformation efficiency was seen, the gDNA was used a second time transforming both the *alrA* null strain and the wild-type strain in parallel to confirm that it was the genetic background and not the quality of the DNA preparation causing the reduced transformation efficiency. Where reduced transformation was confirmed, the resulting colonies were then screened for the presence of expected resistance markers for the respective mutations (zeocine for *alrA*, and erythromycin for pMUTIN,).

### Stabilisation of pLOSS*alrA by deletion of datA

Plasmid pLOSS* *alrA*^*W23*^ was constructed by cloning a PCR generated fragment of DNA corresponding to the *alrA* gene encoded by *B. subtilis* W23 into the pLOSS* vector described by Claessen et al. (35), inserting it such that it was under the control of the P_*spac*_ promoter (using *Bam*HI and *Xba*I sites introduced by the PCR oligonucleotides alrAW23-1 and alrAW23-2). Strain RD180 (*ΔalrA::zeo*) was then transformed with pLOSS*\* alrA*^*W23*^ to give strain KS21 (*ΔalrA* pLOSS*\* alrA*^*W23*^), which exhibited plasmid instability when grown in the presence of D-alanine (see results).

Strain KS21 was then transformed with gDNA from BKE30530 (*ΔdatA::erm*) selecting for erythromycin resistance to produce strain KS23 strain (*ΔdatA ΔalrA* pLOSS*\* alrA*^*W23*^). These two were grown on plates of nutrient agar supplemented with X-gal and with and without D-alanine at 37 °C to determine the stability of the plasmid as indicated by the loss of β-galactosidase activity.

### Complementation of datA

The coding sequence of the *datA* gene and 30 bases upstream, to include the putative ribosome binding site (RBS) of the gene, was inserted into plasmid pPY18 (*bla amyE3’ cat P*_*spac*_ la*cI amyE5’;*(53)) to produce plasmid pKS1 (*bla amyE3’ cat P*_*spac*_ *datA lacI amyE5’*). Plasmid pKS6 was similarly constructed, but in this case the native ribosome binding of *datA* was modified (to a sequence corresponding to the RBS identified upstream of *ftsL*). Both pKS1 and pKS6 were then used to transform strain KS22 (*ΔdatA::erm*) such that the *amyE* locus on the chromosome was replaced by the inducible copy of *datA* to give strains, KS26 and KS41. Correct replacements were confirmed by determining the loss of amylase activity in the strain and by PCR. Both of these strains were then transformed with the gDNA from RD180 strain (*ΔalrA::zeo*) selecting for zeocine resistance in the presence of 0.5 mM D-alanine and 1.0 mM IPTG to give strains KS27 and KS42.

### Cross-feeding assay

In this assay the D-alanine auxotroph strain (RD180) was grown as close as possible to wild type or test strains on a solid medium, ensuring that the strains did not become mixed. Through trial and error it was determined that the D-alanine prototroph (donor; *alrA*^+^) had to be inoculated onto the plate first and incubated for 3.0 h at 37°C prior to inoculating the D-alanine auxotroph (recipient) strain in parallel to the donor strain. The plate was then incubated for a further 23 h and the resulting growth photographed.

### Culture sample processing for RP-HPLC analysis for D-alanine

Samples were taken from the culture at different time points were centrifuged at 3000 × g for 5.0 min and the culture supernatant was filtered through sterile syringe filter (pore size 0.2 μm, GILSON^®^). The resulting filtrate was then passed through a spin column (Vivaspin 2.0 Hydrosart, 2000 MWCO, Generon Ltd.) before analysis. For standards, the LB medium sample was similarly processed with and without added known concentrations of D-alanine. A 150 μl of the “super” filtered culture supernatant was concentrated to approximately 50 μl using a SpeedVac concentrator (SavantSPD131DDA, Thermo Electron Corporation). To this material 150 μl of Marfey’s reagent (1.0 % Nα-(2,4-Dinitro-5-fluorophenyl)-L-alaninamide in acetone) was added and mixed. Then, 40 μl of sodium bicarbonate (1.0 M) was added and mixed prior to incubation for 1.0 h at 40 C° with shaking at 750 rpm (Thermomixer compact, Eppendorf). The mixture was then allowed to cool and 25 μl of HCl (2.2 M) was added to stop the reaction and dried by SpeedVac before being suspended in 150 μl of HPLC suspension solution (90 % of 0.05M triethylamaine phosphate (pH 3.0) and 10 % of acetonitile + 0.1 % formic acid). The resulting suspension was then filtered through 0.22 μm centrifuge tube filter (cellulose acetate filter, Costar) at full speed in a micro centrifuge for 1.0 min, and the filtrate was use for HPLC analysis. Results shown in Fig 5 are representative for 3 independent experiments.

Samples labelled by Marfeys reagent (36) were analysed using a Perkin Elmer series 200 HPLC with a Spheri-5, RP-18 column, dimension 100 mm × 4.6 mm and particle size 5.0 μm (Perkin Elmer) and a column guard, NewGuard RP-18, 7.0 μm, 15 × 3.2 mm (Perkin Elmer). The mobile phase was generated from 0.05 M triethylamine phosphate (pH 3.0) (solution A) and acetonitrile + 0.1 % formic acid (solution B) as a linear gradient of solution B from 10 % to 25 % in 40 min. The flow rate was 1.0 ml/min at 35 °C and the detection wave length was 340 nm, using a S200 Diode array detector (DAD). After each run, the column was washed with water (solution D) and then with 60 % methanol (solution C). The details of the run parameters are shown in (Table S3).

### Determination of amino acid changes in culture media

To identify what effect the *datA* mutation had on the assimilation/release of alanine in a growing culture, parallel cultures of the wild-type (168) and the *datA* null strain (KS22) were grown overnight at 30 °C in a defined minimal medium; GM Opt (M. Chow, personal communication) which was GM media supplemented with amino acids serine, glutamine, asparagine and glutamate at 0.5 mM and all others at 0.25 mM except alanine which was omitted. The cultures were then diluted 1/50 in fresh media and incubated at 37 degrees for 1 h prior to being diluted in the same medium to an optical density of 0.05 at 600 nm. At this point the cultures were divided into 3 aliquots. The first was supplemented with D-alanine at 0.5 mM, and the second L-alanine (0.5 mM) and the last was left without any addition. At regular intervals the culture optical density was determined and 1 ml samples were centrifuged to obtain the culture supernatant, which was filtered through a Microcon centrifugal filter unit (MWCO 3kDa; Sigma) and stored at −20 °C.

The frozen samples were then processed for HPLC analysis to determine the concentration of alanine present using pre-column OPA derivatisation as described by Henderson *et al*. (2008) (54) using an Agilent 1100 HPLC with a Poroshell 120 C18 4 ×m, 4.6 × 150 mm column. The resulting HPLC traces were then analysed with reference to a T0 sample and known standard concentrations of alanine to permit the identification of the peak corresponding to alanine in the trace and to allow its concentration to be determined. The results were then plotted against time and the values obtained for a typical experiment are shown in figure 5.

## Acknowledgments

We would like to thank D. Thwaites for his technical help and advice with respect to amino acid transporters, A. Guyet and L. J. Wu for critical reading of the manuscript. This work was supported by the Kurdistan regional government-IRAQ in the form of a studentship for KS, and Engineering and Physical Sciences Research Council grant EP/N031962/1 (RD)

## Supplemental figures, methods and materials

**Figure S1.**
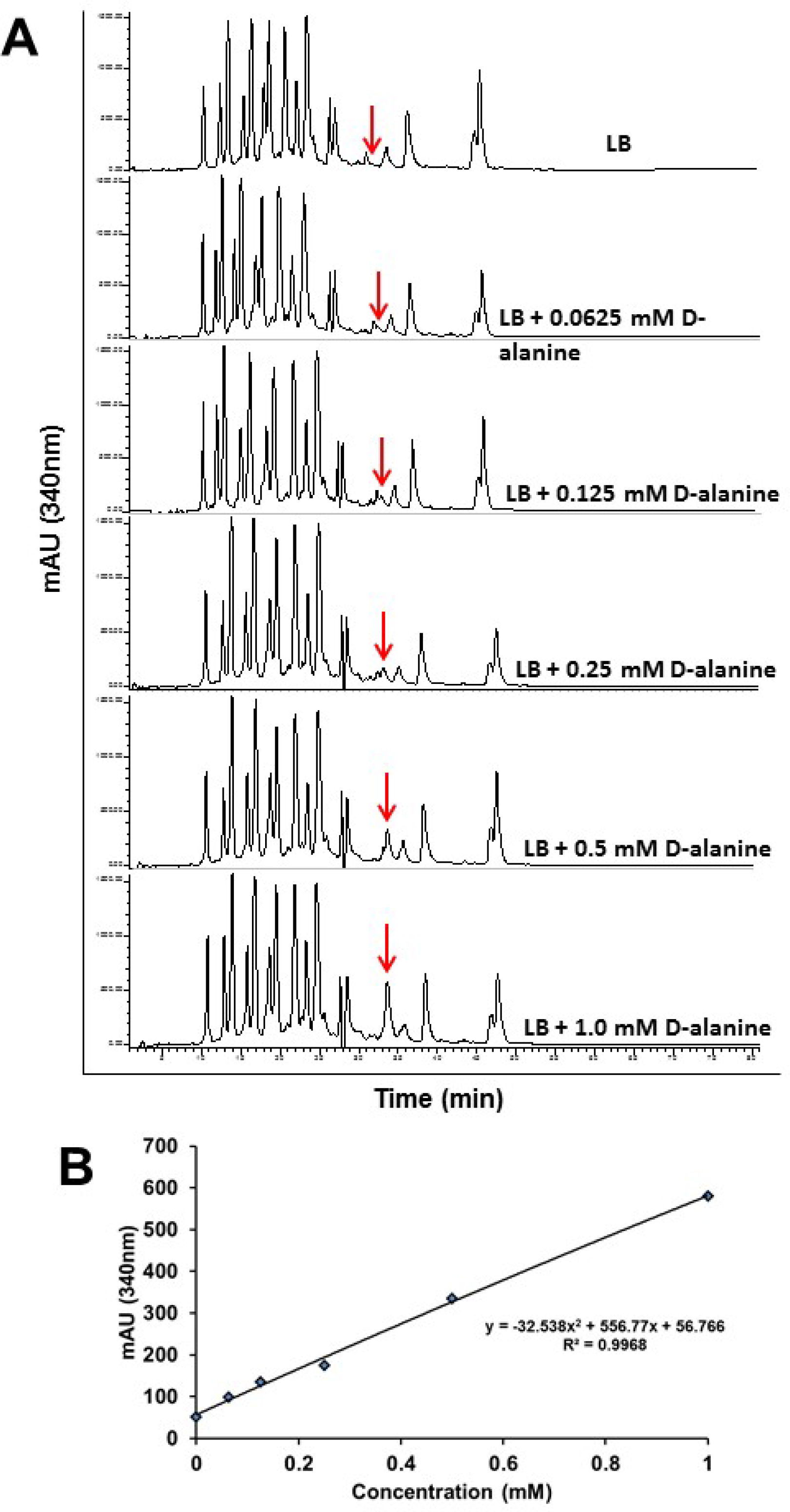
Standard curve for D-alanine quantification in LB media. A) The RP-HPLC analysis of 2000 MWCO filtered LB medium with and without D-alanine (mM). The red arrows indicate the D-alanine peaks, which were raised around 34 min. B) A standard curve was generated from the data of figure (A) for quantitative determination of D-alanine in the growth medium.

**Figure S2.**
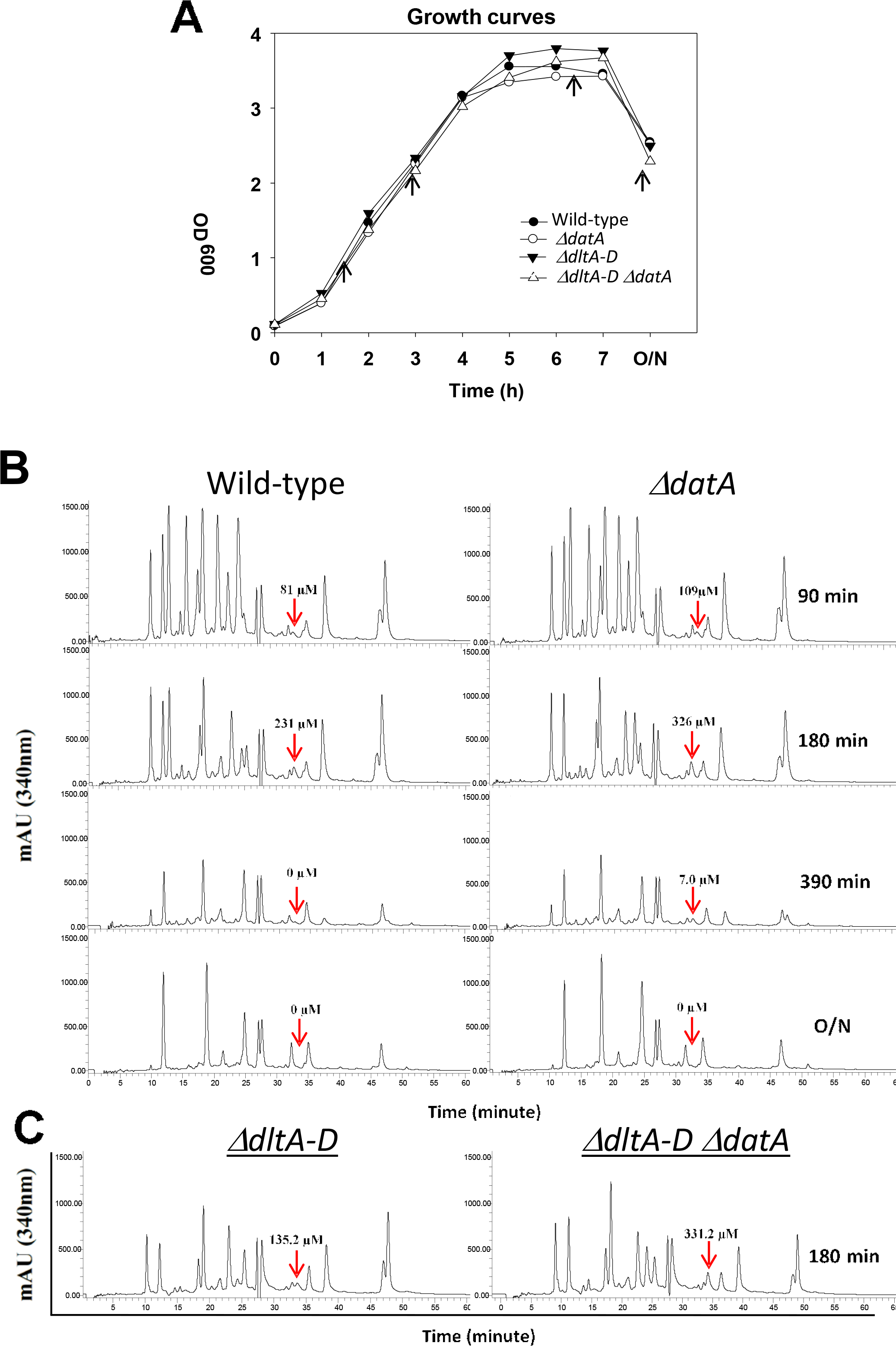
RP-HPLC analysis for the detection of D-alanine in culture supernatants. **Panel A** - Growth curves for cultures samples for HPLC analysis (panel B) the wild-type strain (•) 168CA), *ΔdatA* (○; KS22), *ΔdltA-D* (▾; KS11) and *ΔdltA-D datA* (Δ; KS36) strains in LB medium at 37 °C. The black arrows represent the time points, when the samples were taken for RP-HPLC analysis. **Panel B** - RP-HPLC analysis of 2000 MWCO filtered culture supernatants of 168CA and *ΔdatA* (KS22) strains at different time points (90, 180, 390 min and O/N). **Panal C** - RP-HPLC analysis of *ΔdltA-D* (KS11) and *ΔdltA-D ΔdatA* (KS36) strains at 180 min of incubation. The red arrows indicate D-alanine peaks, which appeared at around 34 min.

**Figure S3.**
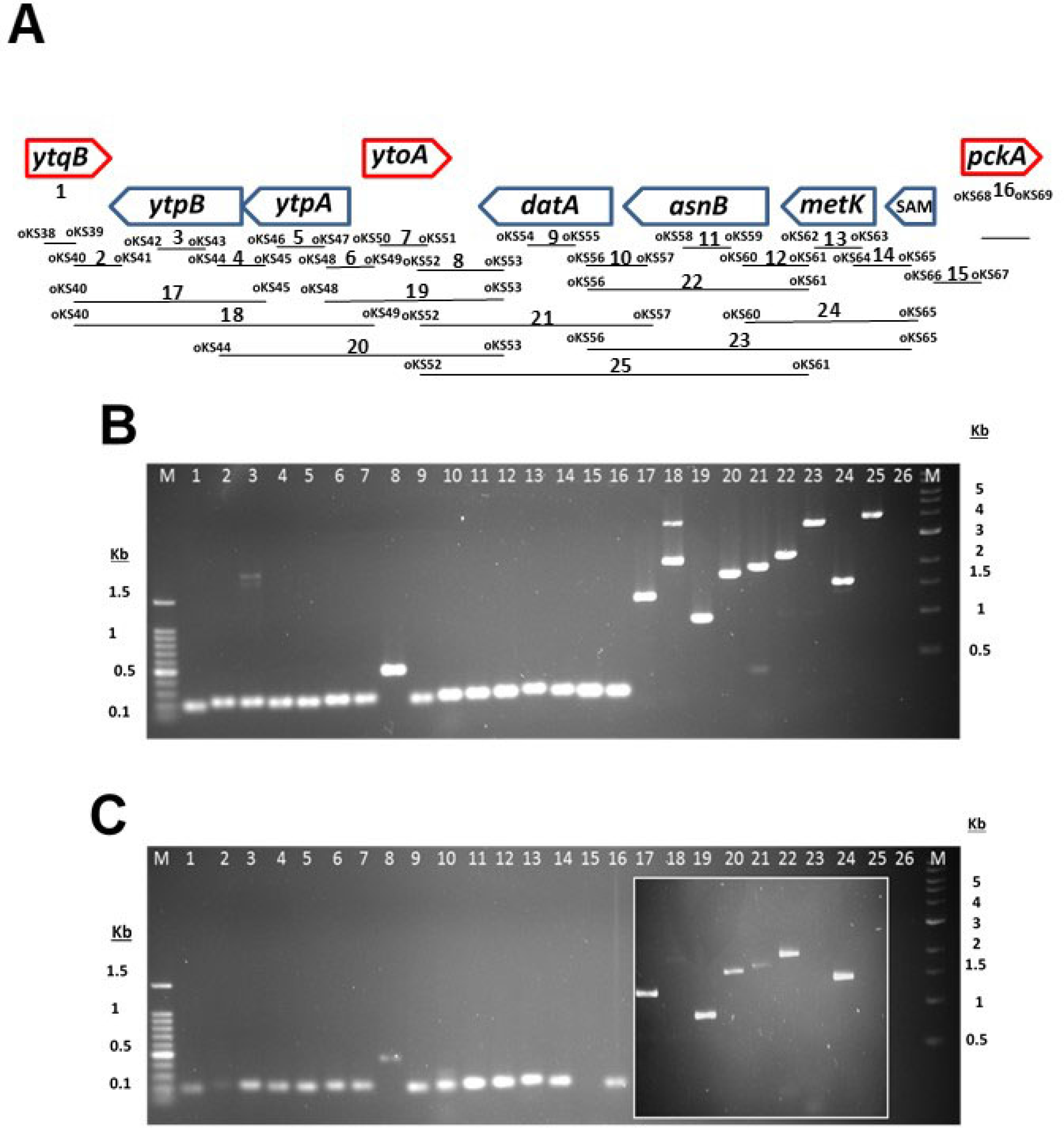
Complementation and Transcription analysis of *datA* gene. **Panel A** - Growth of KS22 (*ΔdatA::erm*), RD180 (*ΔalrA::zeo*) and KS42 (*ΔdatA::erm amyE Ω (cat Pspac datA-ftsL*(RBS)) *ΔalrA::zeo*) strains on nutrient agar (NA) plates with and without 1.0 mM IPTG and 0.5 mM D-alanine. The plates were incubated at 37 °C overnight. **Panel B** provides a diagrammatic representation of the chromosomal region up and downstream of the *datA* gene below which are shown numbered DNA fragments which were amplified by PCR. The numbers indicate which specific pair of oligonucleotides (sequences of which can be found in Table S2) were used to generate the DNA fragment. The images of agarose gel show products of PCR reactions (32 cycles), using gDNA (**panel C**) and cDNA (**panel D**) as templates. The lane numbers correspond to the numbered bars in panel A.

**Figure S4.**
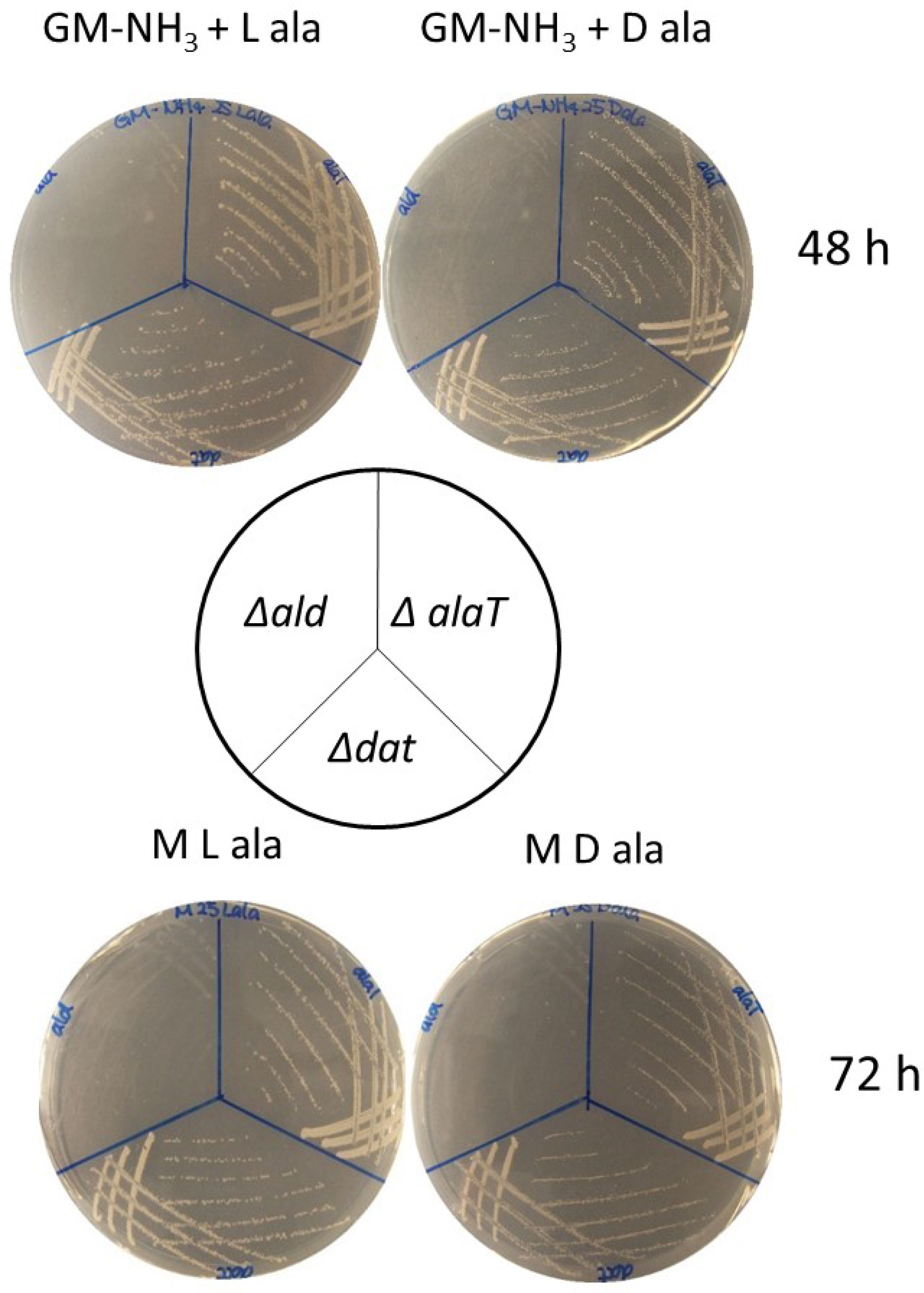
Genes involved in the use of alanine as sole carbon/nitrogen source. For testing alanine used as sole carbon source, *ΔalaT* (MC2) *Δdat* (MC3) and *Δald* (MC5) were grown on M plates with 25 mM D-/L-alanine. For testing alanine used as nitrogen source, these three strains were grown on GM-NH3 plates supplemented with 25 mM D-/L-alanine. The plates were incubated at 37°C for 48 hours.

### Real-time PCR assay

The wild type strain (168CA) was grown in LB media to mid-exponential growth (OD_600_ ~ 1.0) at 37 °C. Total RNA was extracted from 1.0 ml the culture, using total RNA purification plus Kit (NORGEN Biotek Corp.) according to the manufacturer’s protocol. The high capacity cDNA reverse transcription kit (Applied Biosystems^TM^) was used for making cDNA according to manufacturer’s protocol. The total RNA extract was used as template and non-specific random hexamers as primer in cDNA production. Real-time PCR was then performed on normal PCR machine, using DNA as template, gene specific primers and Q5^TM^ High-Fidelity DNA polymerase (New England BioLabs®) according to the manufacturer’s instructions. The Real-time PCR products were run on agarose gel (Fig S2).

**Table S1.** Plasmid constructs used in this study.

| Name plasmid | Relevant features | Construction | Source/Reference |
| --- | --- | --- | --- |
| pPY18 | *bla amyE3' cat P<sub>spac</sub> lacI amyE5'* | ----- | (53) |
| pSG1 | *bla cat* | ----- | (55) |
| pLOSS* *alrA* | *bla spc P<sub>spac</sub> alrA<sup>W23</sup> - PdivlA- lacZ lacI rep pLS20\** | Insertion of *alrA* gene into *pLOSS\** vector | This work |
| pKS1 | *bla amyE3' cat P<sub>spac</sub> datA*<br>*lacI amyE5'* | Insertion of *datA* gene and its upstream 30 bases into *Xba*I cut pPY18 | This work |
| pKS6 | *bla amyE3' cat P<sub>spac</sub> datA-ftsL(RBS) lacI lacZ amyE5'* | Insertion of *datA* gene and RBS of *ftsL* gene into *Xba*I and *Bgl*II cut pPY18 | This work |

**Table S2.**
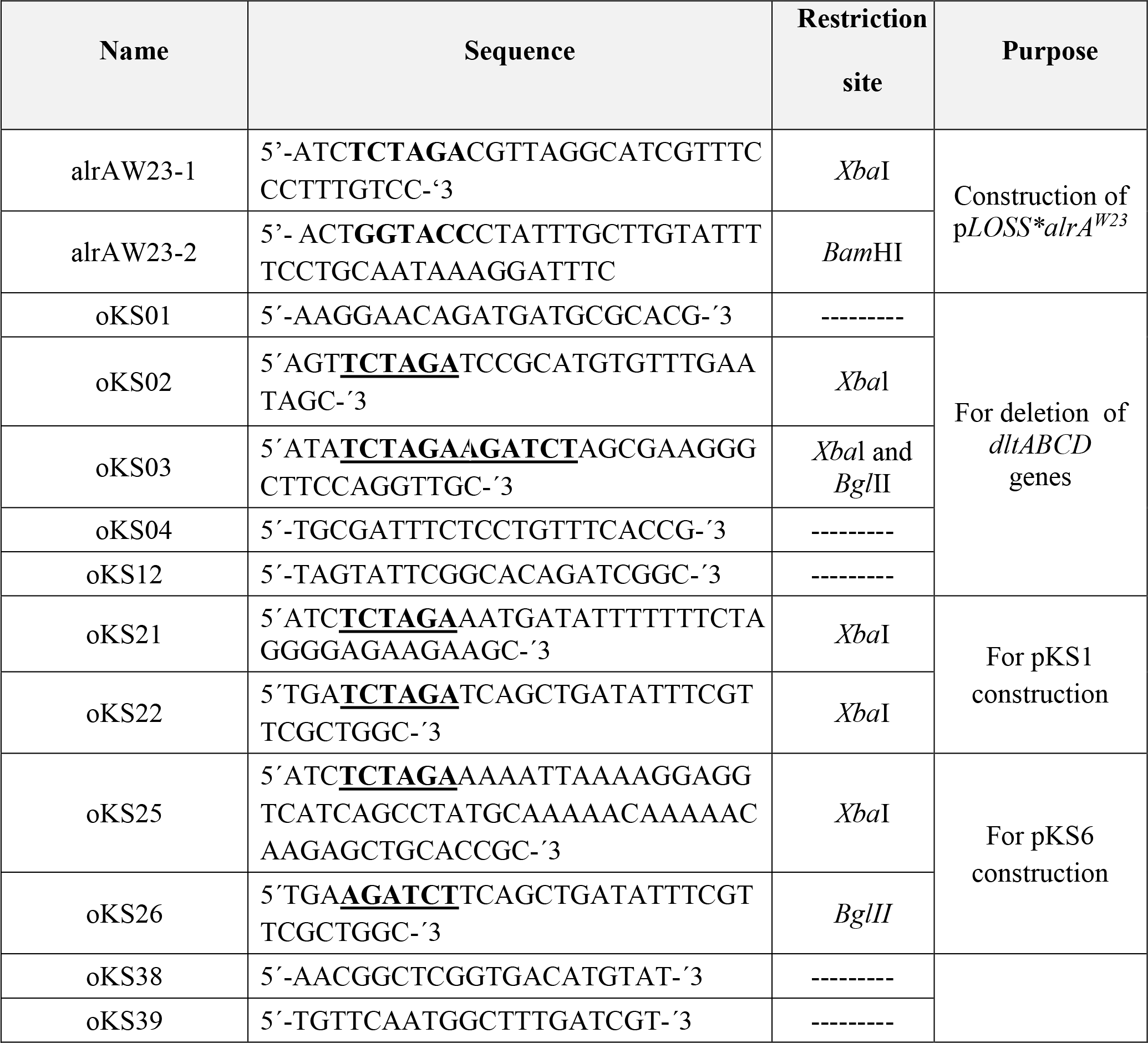
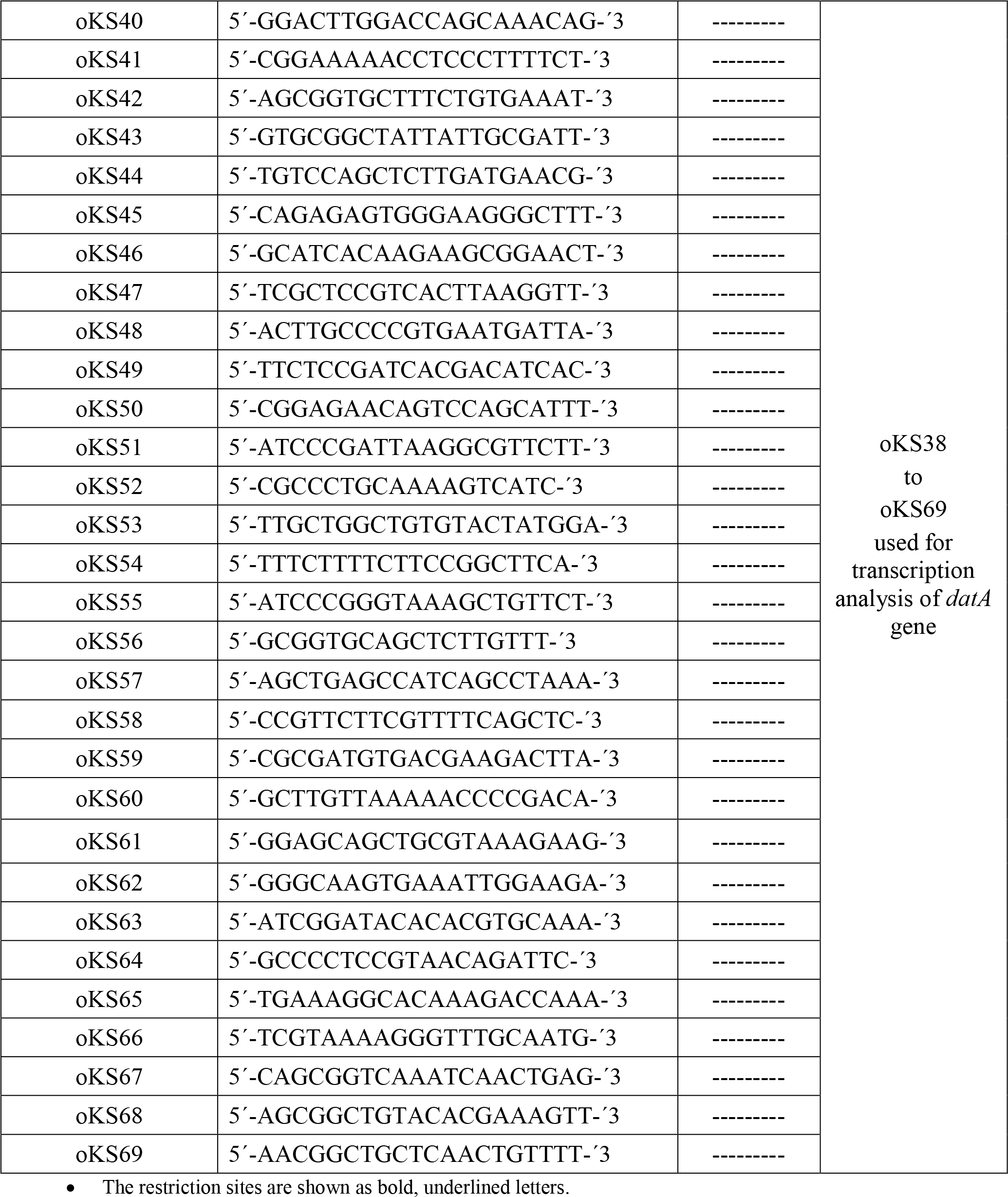
Oligonucleotides.

**Table S3.** Pump and run parameters of RP-HPLC analysis.

| Step | Time (min) | Flow (ml/min) | A | B | C | D | Curve | Sample volume(μl) |
| --- | --- | --- | --- | --- | --- | --- | --- | --- |
| 0 | 0.5 | 1.00 | 90.0 | 10.0 | 0.0 | 0.0 | 0.0 | 150 |
| 1 | 1.5 | 1.00 | 90.0 | 10.0 | 0.0 | 0.0 | 0.0 |  |
| 2 | 40.0 | 1.00 | 75.0 | 25.0 | 0.0 | 0.0 | 1.0 |  |
| 3 | 25.0 | 1.00 | 60.0 | 40.0 | 0.0 | 0.0 | 1.0 |  |
| 4 | 0.5 | 1.00 | 90.0 | 10.0 | 0.0 | 0.0 | 0.0 |  |
| 5 | 20.0 | 1.00 | 90.0 | 10.0 | 0.0 | 0.0 | 0.0 |  |
| Total Run Time | 87.00 |  |  |  |  |  |  |  |

**Table S4.**
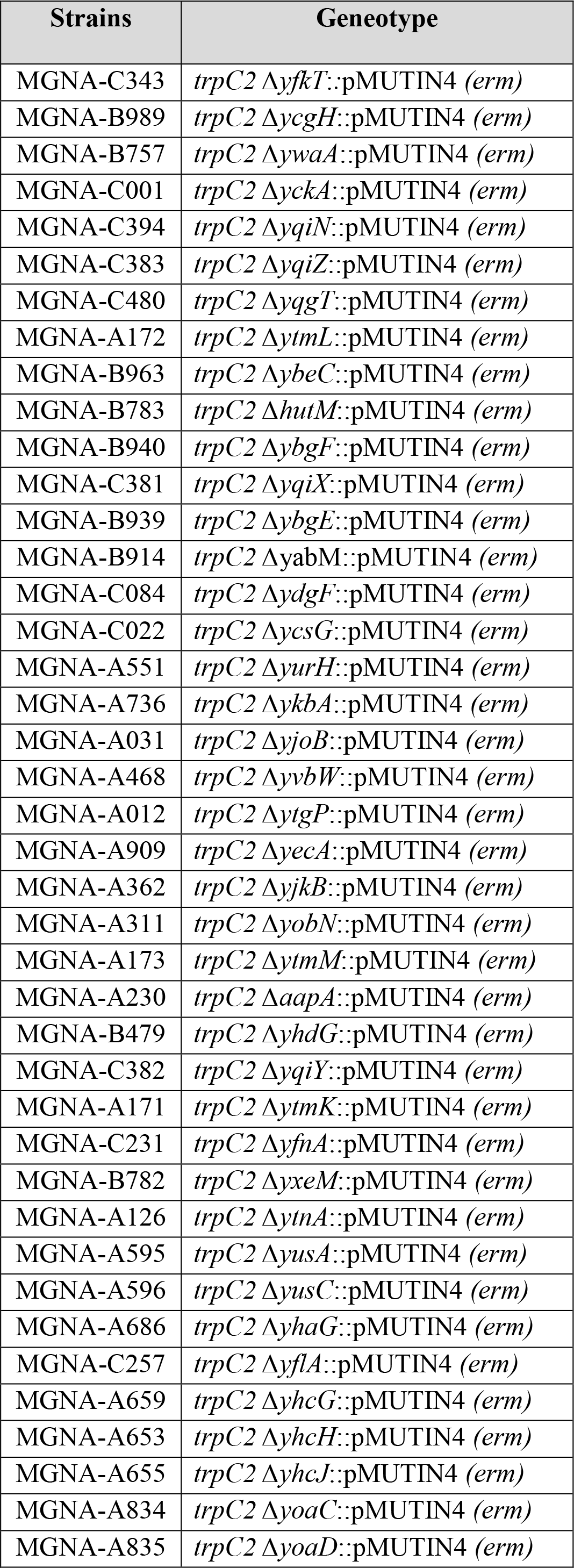
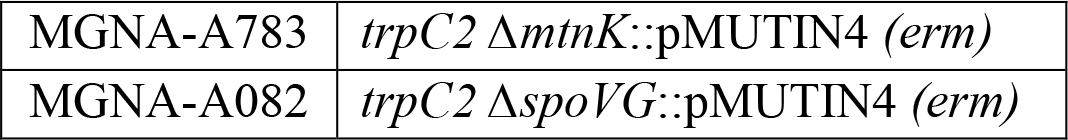
Putative amino acids transport mutants.

## References

1. Lam H, Oh DC, Cava F, Takacs CN, Clardy J, de Pedro MA, et al. D-amino acids govern stationary phase cell wall remodeling in bacteria. Science. 2009;325(5947):1552–5.

2. Hernandez SB, Cava F. Environmental roles of microbial amino acid racemases. Environmental Microbiology. 2016;18(6):1673–85.

3. Perego M, Glaser P, Minutello A, Strauch MA, Leopold K, Fischer W. Incorporation of D-alanine into lipoteichoic acid and wall teichoic acid in *Bacillus subtilis*. Identification of genes and regulation. The Journal of Biological Chemistry. 1995;270(26):15598–606.

4. Wecke J, Perego M, Fischer W, Perego M, Glaser P, Minutello A, et al. D-alanine deprivation of *Bacillus subtilis* teichoic acids is without effect on cell growth and morphology but affects the autolytic activity Incorporation of D-alanine into lipoteichoic acid and wall teichoic acid in *Bacillus subtilis*. Identification of genes and regulation. Microb Drug Resist. 1996;2(1):123–9.

5. Peng B, Su YB, Li H, Han Y, Guo C, Tian YM, et al. Exogenous alanine and/or glucose plus kanamycin kills antibiotic-resistant bacteria. Cell Metabolism. 2015;21(2):249–62.

6. Yasuda Y, Tochikubo K. Relation between D-glucose and L- and D-alanine in the initiation of germination of Bacillus subtilis spore. Microbiology and immunology. 1984;28(2):197–207.

7. Shrestha R, Lockless SW, Sorg JA. A *Clostridium difficile* alanine racemase affects spore germination and accommodates serine as a substrate. The Journal of Biological Chemistry. 2017;292(25):10735–42.

8. McKevitt MT, Bryant KM, Shakir SM, Larabee JL, Blanke SR, Lovchik J, et al. Effects of endogenous D-alanine synthesis and autoinhibition of *Bacillus anthracis* germination on in vitro and in vivo infections. Infection and Immunity. 2007;75(12):5726–34.

9. Ferrari E, Henner DJ, Yang MY. Isolation of an Alanine Racemase Gene from *Bacillus subtilis* and its Use for Plasmid Maintenance in *B. subtilis*. Bio/Technology. 1985;3:1003.

10. Xia Y, Chen W, Zhao J, Tian F, Zhang H, Ding X. Construction of a new food-grade expression system for *Bacillus subtilis* based on theta replication plasmids and auxotrophic complementation. Applied Microbiology and Biotechnology. 2007;76(3):643–50.

11. Koch AL, Doyle RJ. Inside-to-outside growth and turnover of the wall of Gram-positive rods. Journal of theoretical biology. 1985;117(1):137–57.

12. Pooley HM. Layered distribution, according to age, within the cell wall of *Bacillus subtilis*. Journal of Bacteriology. 1976;125(3):1139–47.

13. Pooley HM. Turnover and spreading of old wall during surface growth of *Bacillus subtilis*. Journal of Bacteriology. 1976;125(3):1127–38.

14. Atrih A, Bacher G, Allmaier G, Williamson MP, Foster SJ. Analysis of peptidoglycan structure from vegetative cells of *Bacillus subtilis* 168 and role of PBP 5 in peptidoglycan maturation. Journal of Bacteriology. 1999;181(13):3956–66.

15. Hoyland CN, Aldridge C, Cleverley RM, Duchene MC, Minasov G, Onopriyenko O, et al. Structure of the LdcB LD-carboxypeptidase reveals the molecular basis of peptidoglycan recognition. Structure. 2014;22(7):949–60.

16. Neuhaus FC, Baddiley J. A continuum of anionic charge: structures and functions of D-alanyl-teichoic acids in Gram-positive bacteria. Microbiol Mol Biol Rev. 2003;67(4):686–723.

17. Radkov AD, Moe LA. Bacterial synthesis of D-amino acids. Applied Microbiology and Biotechnology. 2014;98(12):5363–74.

18. Walsh CT. Enzymes in the D-alanine branch of bacterial cell wall peptidoglycan assembly. The Journal of Biological Chemistry. 1989;264(5):2393–6.

19. Pierce KJ, Salifu SP, Tangney M. Gene cloning and characterization of a second alanine racemase from *Bacillus subtilis* encoded by yncD. FEMS microbiology letters. 2008;283(1):69–74.

20. Belitsky BR. Biosynthesis of Amino acids of the Glutamate and Aspartate Families, Alanine, and Polyamines. In: Sonenshein AL, editor. Bacillus subtilis and its Closest Relatives: from Genes to cells Washington, D. C.: ASM Press; 2002. p. 203–31.

21. Fotheringham IG, Bledig SA, Taylor PP. Characterization of the genes encoding D-amino acid transaminase and glutamate; racemase, two D-glutamate biosynthetic enzymes of Bacillus sphaericus ATCC 10208. Journal of bacteriology. 1998;180(16):4319–23.

22. Berberich R, Kaback M, Freese E, Reese I. D-amino acids as inducers of L-alanine dehydrogenase in *Bacillus subtilis*. J Biol Chem. 1968;243:1006–11.

23. Freese E, Park SW, Cashel M. The Developmental significance of alanine dehydrogenase in *Bacillus subtilis*. Proceedings of the National Academy of Sciences of the United States of America. 1964;51:1164–72.

24. Siranosian KJ, Ireton K, Grossman AD. Alanine dehydrogenase (ald) is required for normal sporulation in *Bacillus subtilis*. Journal of Bacteriology. 1993;175(21):6789–96.

25. Stannek L, Thiele MJ, Ischebeck T, Gunka K, Hammer E, Volker U, et al. Evidence for synergistic control of glutamate biosynthesis by glutamate dehydrogenases and glutamate in *Bacillus subtilis*. Environmental Microbiology. 2015;17(9):3379–90.

26. Osborne SE, Tuinema BR, Mok MC, Lau PS, Bui NK, Tomljenovic-Berube AM, et al. Characterization of DalS, an ATP-binding cassette transporter for D-alanine, and its role in pathogenesis in Salmonella enterica. The Journal of Biological Chemistry. 2012;287(19):15242–50.

27. Yu J, Ge J, Heuveling J, Schneider E, Yang M. Structural basis for substrate specificity of an amino acid ABC transporter. Proceedings of the National Academy of Sciences of the United States of America. 2015;112:5243–8.

28. M. J Zimbro, D. A. Power, S. M. Miller, G. E. Wilson, J. A. Johnson. Difco⍰& BBL⍰ Manual Manual of Microbiological Culture Media. In: Becton DaC, Sparks MU, editors. 2nd ed: BD Diagnostics – Diagnostic Systems; 2009.

29. Steil L, Serrano M, Henriques AO, Volker U. Genome-wide analysis of temporally regulated and compartment-specific gene expression in sporulating cells of *Bacillus subtilis*. Microbiology. 2005;151(Pt 2):399–420.

30. Kimura K, Tran LS, Itoh Y. Roles and regulation of the glutamate racemase isogenes, *racE* and *yrpC*, in *Bacillus subtilis*. Microbiology. 2004;150(Pt 9):2911–20.

31. Miyamoto T, Katane M, Saitoh Y, Sekine M, Homma H. Identification and characterization of novel broad-spectrum amino acid racemases from *Escherichia coli* and *Bacillus subtilis*. Amino Acids. 2017;49(11):1885–94.

32. Thompson RJ, Bouwer HG, Portnoy DA, Frankel FR. Pathogenicity and immunogenicity of a *Listeria monocytogenes* strain that requires D-alanine for growth. Infection and Immunity. 1998;66(8):3552–61.

33. Mortuza R, Aung HL, Taiaroa G, Opel-Reading HK, Kleffmann T, Cook GM, et al. Overexpression of a newly identified D-amino acid transaminase in *Mycobacterium smegmatis* complements glutamate racemase deletion. Molecular Microbiology. 2018;107:198–213.

34. Vagner V, Dervyn E, Ehrlich SD. A vector for systematic gene inactivation in *Bacillus subtilis*. Microbiology. 1998;144 (Pt 11):3097–104.

35. Claessen D, Emmins R, Hamoen LW, Daniel RA, Errington J, Edwards DH. Control of the cell elongation-division cycle by shuttling of PBP1 protein in *Bacillus subtilis*. Molecular Microbiology. 2008;68(4):1029–46.

36. Szokan G, Mezo G, Hudecz F. Application of Marfey’s reagent in racemization studies of amino acids and peptides. J Chromatogr. 1988;444:115–22.

37. Nicolas P, Mader U, Dervyn E, Rochat T, Leduc A, Pigeonneau N, et al. Condition-dependent transcriptome reveals high-level regulatory architecture in *Bacillus subtilis*. Science. 2012;335(6072):1103–6.

38. Dajkovic A, Tesson B, Chauhan S, Courtin P, Keary R, Flores P, et al. Hydrolysis of peptidoglycan is modulated by amidation of meso-diaminopimelic acid and Mg(2+) in *Bacillus subtilis*. Molecular Microbiology. 2017;104(6):972–88.

39. Rigden DJ, Fernández XM. The 2018 Nucleic Acids Research database issue and the online molecular biology database collection. Nucleic Acids Research. 2018;46(D1):D1–D7.

40. Yoneyama H, Hori H, Lim SJ, Murata T, Ando T, Isogai E, et al. Isolation of a mutant auxotrophic for L-alanine and identification of three major aminotransferases that synthesize L-alanine in *Escherichia coli*. Biosci Biotechnol Biochem. 2011;75(5):930–8.

41. Tam le T, Eymann C, Antelmann H, Albrecht D, Hecker M. Global gene expression profiling of *Bacillus subtilis* in response to ammonium and tryptophan starvation as revealed by transcriptome and proteome analysis. Journal of molecular microbiology and biotechnology. 2007;12(1-2):121–30.

42. Siranosian KJ, Ireton K, Grossman AD. Alanine dehydrogenase (*ald*) is required for normal sporulation in *Bacillus subtilis*. Journal of Bacteriology. 1993;175:6789–96.

43. Borisova M, Gaupp R, Duckworth A, Schneider A, Dalugge D, Muhleck M, et al. Peptidoglycan Recycling in Gram-Positive Bacteria Is Crucial for Survival in Stationary Phase. mBio. 2016;7(5).

44. Reith J, Mayer C. Peptidoglycan turnover and recycling in Gram-positive bacteria. Applied microbiology and biotechnology. 2011;92(1):1–11.

45. Taylor PP, Fotheringham IG. Nucleotide sequence of the *Bacillus licheniformis* ATCC 10716 dat gene and comparison of the predicted amino acid sequence with those of other bacterial species. Biochim Biophys Acta. 1997;1350(1):38–40.

46. Martinez del pozo A, Merola M, Ueno H, Manning JM, Tanizawa K, Nishimura K, et al. Activity and spectroscopic properties of bacterial D-amino acid transaminase after multiple site-directed mutagenesis of a single tryptophan residue. Biochemistry. 1989;28(2):510–6.

47. Ward JB, Zahler SA. Genetic studies of leucine biosynthesis in *Bacillus subtilis*. Journal of Bacteriology. 1973;116:719–26.

48. Anagnostopoulos C, Spizizen J. Requirements for transformation in *Bacillus subtilis*. Journal of Bacteriology. 1961;81(5):741–6.

49. Young FE, Spizizen J. Physiological and genetic factors affecting transformation of *Bacillus subtilis*. Journal of Bacteriology. 1961;81:823–9.

50. D H. Techniques for transformation of E.coli. In: Glover DMIP, editor. DNA cloning - A practical approach Oxford. Washington D.C: pp. 109–135. 1985.

51. Koo BM, Kritikos G, Farelli JD, Todor H, Tong K, Kimsey H, et al. Construction and Analysis of Two Genome-Scale Deletion Libraries for *Bacillus subtilis*. Cell Syst. 2017;4(3):291–305.e7.

52. Kobayashi K, Ehrlich SD, Albertini A, Amati G, Andersen KK, Arnaud M, et al. Essential *Bacillus subtilis* genes. Proceedings of the National Academy of Sciences of the United States of America. 2003;100(8):4678–83.

53. Duggin IG, Andersen PA, Smith MT, Wilce JA, King GF, Wake RG. Site-directed mutants of RTP of *Bacillus subtilis* and the mechanism of replication fork arrest. J Mol Biol. 1999;286(5):1325–35.

54. Henderson JWR, R. D.; Bidlingmeyer, B. A.; Woodward, C. Rapid, Accurate, Sensitive, and Reproducible HPLC Analysis of Amino Acids. Agilent Technical note. 2008.

55. Errington J. Efficient *Bacillus subtilis* cloning system using bacteriophage vector phi 105J9. J Gen Microbiol. 1984;130(10):2615–28.

